# Host network-based discovery of critical regulators of innate immunity, virus growth, and pathogenesis in influenza virus infection

**DOI:** 10.1101/2022.08.29.505723

**Authors:** Amie J. Eisfeld, Shufang Fan, Hongyu Rao, Backiyalakshmi Ammayappan Venkatachalam, Danielle Westhoff Smith, Peter J. Halfmann, Kevin B. Walters, Sharmila Nair, Larissa B. Thackray, Jacob F. Kocher, Amy C. Sims, Hugh D. Mitchell, Gabriele Neumann, Volker Blank, Katrina M. Waters, Ralph S. Baric, Michael S. Diamond, Yoshihiro Kawaoka

## Abstract

Innate immunity is protective against viruses, but also can facilitate pathological infection responses. Despite intensive research, our understanding of the mechanisms that regulate innate immunity in virus infection remains incomplete. Systems biology-based data-driven modeling approaches hold substantial promise toward discovery of crucial innate immune signaling regulators, yet model-derived predictions are almost completely unexplored. Here, we carried out systematic experimental validation of candidate regulators predicted by a transcriptional association network model of influenza virus-infected cells. We identified dozens of novel innate immune signaling regulators with potent effects on the replication of influenza and other viruses, and importantly, we established the biological relevance of a validated regulator *in vivo*. Collectively, these findings aid in clarifying mechanisms of influenza virus pathogenicity and might lead to innovative approaches for treating influenza virus disease. Similar data-driven modeling strategies may be applicable for the study of other pathogen systems or immunological disorders.

## INTRODUCTION

Influenza viruses pose an ongoing threat to global public health, causing up to 5 million cases of severe disease and 0.65 million deaths annually^1^. Unpredictable but recurrent pandemics are typically associated with higher morbidity and mortality^2^, and sporadic zoonotic infections with high fatality rates occur in laboratory-confirmed cases (*e.g.*, 56% and 39% for highly pathogenic H5N1 and H7N9 avian virus strains, respectively)^3^. Currently, influenza management with available countermeasures is challenging due to sub-optimal vaccine efficacy, vaccine mismatches, and development of antiviral resistance^4, 5^. The discovery of novel host regulators of virus growth and inflammation, which presumably are less susceptible to the selective pressures that give rise to drug resistance, may facilitate host-targeted intervention strategies to curb human morbidity and mortality in future epidemics, pandemics, and zoonotic outbreaks.

Cell autonomous innate immune signaling (*i.e.*, the intrinsic antiviral and pro-inflammatory signaling pathways activated by virus infection in all cell types) limits virus growth and spread in respiratory epithelium at the initial site of influenza virus infection and generates inflammatory signals that communicate infection to the immune system^6, 7^. Pattern recognition receptors (PRRs), including Toll-like receptors (TLRs; primarily TLR3 in respiratory epithelium) and DDX58 (a RIG-I-like receptor), detect influenza virus RNA and trigger production of type I and type III interferons (IFNs), cytokines, and other pro-inflammatory mediators via the IRF3, IRF7, and NFκB transcription factors^8^. Subsequently, secreted IFNs bind to their receptors to mediate autocrine and paracrine signaling and activate expression of hundreds of IFN-stimulated genes (ISGs) via STAT1, STAT2, and IRF9 transcription factors. ISG-encoded proteins that directly interfere with influenza virus replication in human cells include MX1, PKR, TRIM22, IFITM1, IFITM2, IFITM3, ISG15, OAS3, RSAD2 (also known as viperin), BST2 (also known as tetherin), and ISG20, among others^8^. Other influenza virus-associated stimuli activate a third PRR—NLRP3, a NOD-like receptor—leading to IL1B and IL18 cleavage and secretion^9^. Cytokines, chemokines, and pro-inflammatory mediators stimulated by all three PRR pathways promote leukocyte recruitment, proliferation, and differentiation in the lung, aiding viral clearance, tissue repair, and development of long-term immunity^6, 7, 9^. PRR-dependent signaling also may activate programmed cell death, presumably as a mechanism to prevent virus spreading^9–13^, and excessive or prolonged PRR activation may substantially elevate pro-inflammatory cytokine levels, inducing cell death in both infected and bystander cells and contributing to immunopathology and fatal outcomes^11, 14, 15^. While much is known about the relationships between cell autonomous innate immunity, virus growth, and mechanisms of viral pathogenesis, gaps in knowledge still exist, and additional regulators of antiviral and pro-inflammatory signaling have yet to be discovered.

Data-driven modeling, which uses mathematical analyses and computational algorithms to infer biological relationships within large-scale ‘omics profiling data, is a powerful tool for unbiased prediction of candidate regulators of immunological responses^16, 17^, and coupled with experimental validation, has potential to reveal novel or unexpected biological insights for diverse phenotypes in various model systems. For example, Amit *et al.*^18^ used a data-driven transcriptional regulatory network model combined with RNA interference (RNAi) perturbations to predict and validate regulatory functions of >100 transcription factors, chromatin modifiers, and RNA binding proteins in pathogen-sensing pathways in human dendritic cells. We^19–22^ and others^23–25^ have used data-driven modeling to evaluate influenza virus-induced host responses and predict candidate regulators of virus replication, cell autonomous innate immune signaling, or influenza pathogenicity. However, only a few model-based candidate regulator predictions have been tested for effect(s) on influenza infection-associated phenotype(s)^19, 20, 22, 23^, and none of the models has been validated extensively at the experimental level.

Previously, we established a host response transcriptional association network model of influenza virus-infected cells, devised a network-based ranking strategy to identify candidate regulators of influenza virus growth and pathogenesis, and validated our ranking strategy by using computational methods^21^. Here, we carried out systematic experimental validation of top regulator candidates predicted by this approach, most of which have no well-established role in innate immunity in viral infections. We found that a high proportion of candidate host gene regulators exhibit antiviral activity against influenza virus, and that most of the antiviral factors also regulate cell autonomous innate immune signaling. We also showed that a subset could regulate growth of representative viruses from *Coronaviridae*, *Filoviridae*, and/or *Flaviviridae* families. For one antiviral factor (NFE2L3), we demonstrated its ability to limit influenza disease pathogenicity in mice. Our results indicate that candidate host gene regulators predicted from our host transcriptional association network model affect virus growth and disease pathogenesis and may possess conserved activity in response to infection with diverse viruses.

## RESULTS

### Host network model and candidate regulator prioritization

In a previous study^21^, we inferred a host transcriptional association network model from genome-wide transcriptional quantification data of human bronchial epithelial cells responding to infection by either a 2009 pandemic H1N1 influenza virus (pH1N1) or a highly pathogenic H5N1 avian influenza virus (H5N1). The inferred network comprised 11,588 transcripts, each represented by a node, with edges between nodes representing the pairwise expression relationship between transcripts (**Fig. 1a**) and the cumulative relationships of all nodes and edges giving rise to the network’s global configuration (*i.e.*, its ‘topology’). Two important topological components within the network are ‘hub nodes’, which have many associated edges (*i.e.*, are highly connected to other transcripts), and ‘bottleneck nodes’, which have a high number of shortest paths between pairs of nodes passing through them (*i.e.*, they connect large groups of transcripts in different network regions)^26^ (**Fig. 1a**). Both node types represent critical points in the network; specifically, hub node perturbation may disrupt expression of large groups of associated transcripts, whereas bottleneck node perturbation may isolate linked network regions and prevent information flow.

**Figure 1.**
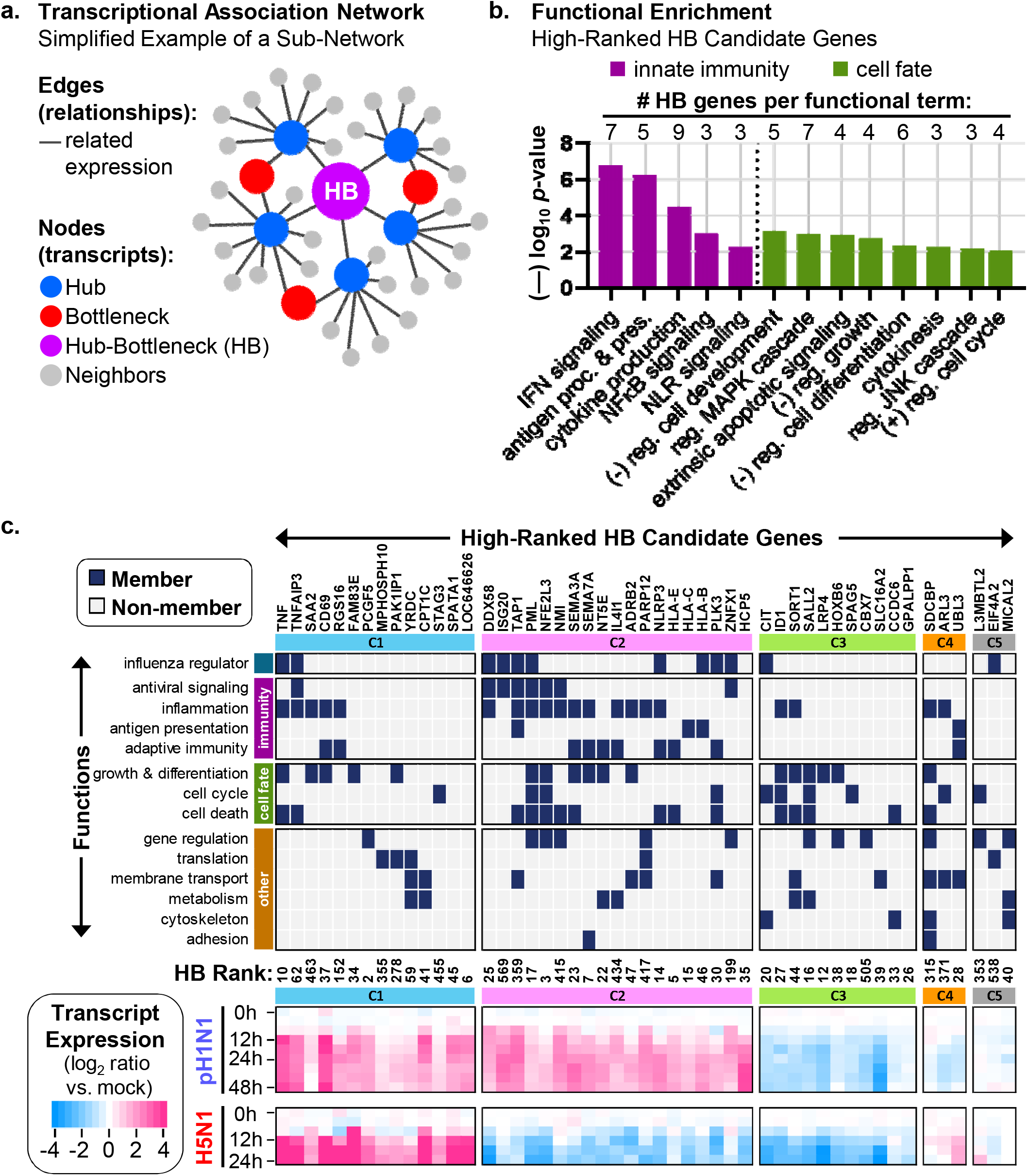
Attributes of high-ranked HB genes. **(a)** The panel depicts spatial organization of a simple transcriptional association network example, including ‘hub’ nodes (with a high number of associated edges), ‘bottleneck’ nodes (with a high number of shortest paths between all pairs of nodes passing through them), and ‘HB nodes’ (with both hub-like and bottleneck-like characteristics). (**b**) For 50 high-ranked HB genes (**Supplementary Table 1a**), the graph shows representative enriched functional terms and enrichment scores (—log10 p-values) determined by the web-based Metascape software (also see **Supplementary Table 1c**). The number of genes comprising each enriched functional term is indicated above each bar on the graph. (**c**) The panel shows individual functions of high-ranked HB genes (if known) (**top**) and their expression in Calu-3 cells infected with pH1N1 (A/California/04/2009) or H5N1 (A/Vietnam/1203/2004) (**bottom**) (also see **Supplementary Table 1e**). HB genes were assigned to expression clusters (C1, C2, C3, C4, or C5), defined by trajectories of expression (significant upregulation or downregulation) in pH1N1 and H5N1 infections, and cluster membership is indicated by the colored bars above the function and expression sub-panels. HB gene ranks (**Supplementary Table 1a**) are shown between the function and expression sub-panels, and a heat map key is given at the bottom left.

We previously hypothesized that hub nodes and/or bottleneck nodes would comprise transcripts derived from genes important for regulating transcriptional programs activated by influenza virus infection, including cell autonomous innate immune signaling, and therefore would impact influenza infection outcomes. To examine this possibility, we ranked nodes by degree scores (for hub-like activity), betweenness scores (for bottleneck-like activity), and a combined degree-betweenness score; and for nodes at the top of each ranked list, we determined enrichment for known regulators of influenza virus replication or pathogenicity^21^. Indeed, we found that nodes with high rankings in both hub-like and bottleneck-like topology (*i.e.*, ‘hub-bottleneck nodes’, hereafter referred to as ‘HB nodes’, ‘HB genes’, or ‘HB candidate genes’) (**Fig. 1a**) are highly enriched for influenza infection regulators^21^. Based on these observations, we further hypothesized that host factors that have not been reported to regulate influenza virus growth and/or cell autonomous innate immunity may be enriched among high-ranked HB nodes. Here, we tested this hypothesis experimentally by focusing on 50 high-ranked HB genes (representing the 0.01—4.91 percentiles, or HB ranks 2—569, of all network nodes; we did not include the top-ranked HB candidate gene, RPSAP44, a pseudogene, due to the lack of available perturbation reagents) (**Supplementary Table 1a**), and for comparison, 22 low-ranked HB genes (representing the 99.28—99.99 percentiles, or HB ranks 11505—11587, of all network nodes) (**Supplementary Table 1b**), which served as controls.

### High-ranked HB genes possess attributes consistent with wide-ranging regulatory roles in influenza virus infection

To better understand how HB genes may regulate influenza infection and pathogenesis, we used Metascape^27^ to identify established functions of high- and low-ranked HB gene sets. High-ranked HB genes are enriched (*p* < 0.01) for pathway, biological process, or gene set terms related to innate immunity or cell fate determination (**Fig. 1b** and **Supplementary Table 1c**; 29 of the 50 high-ranked HB genes are associated with at least one term), whereas low-ranked HB genes exhibit minimal functional enrichment (**Supplementary Fig. 1a** and **Supplementary Table 1d**). For high-ranked HB genes, enriched terms include pathways and processes associated with PRR activation (‘IFN signaling’, ‘NLR signaling’, and ‘NFκB signaling’), control of influenza virus growth (‘regulation of MAPK cascade’ and ‘cell cycle’)^28, 29^, or regulation of autophagy and/or apoptosis (‘regulation of JNK cascade’ and ‘extrinsic apoptotic pathway’)^11, 30, 31^ in influenza virus-infected cells. These observations indicate that a portion of high-ranked HB genes has strong links to—and may regulate—cellular signaling networks with important pro-viral, antiviral, or pro-inflammatory functions. Other high-ranked HB genes lacking association with enriched functional terms may possess unknown connections to the same signaling networks or have other unappreciated roles in viral infection.

To expand on this analysis, we manually integrated enrichment analyses with detailed literature mining to assign specific functions to individual HB genes and grouped them by their overall transcript expression trends in pH1N1 and H5N1 infected cells (**Fig. 1c** and **Supplementary Fig. 1b;** log_2_ fold-change and false discovery rate-adjusted q-values for HB gene transcripts are provided in **Supplementary Tables 1e** and **1f**). Among the high-ranked HB genes are seven known regulators of cell autonomous innate immune signaling in influenza virus-infected epithelial cells, all of which also affect virus replication and/or pathogenicity (*DDX58*, *NLRP3*, *NMI*, *TNF*, *TNFAIP3*, *TAP1*, and *ZNFX1*)^6, 7, 9, 32–39^, and six other genes previously implicated in the regulation of influenza virus growth (*HLA-B*, *PLK3*, *CIT*, *EIF4A2*, *ISG20* and *PML*)^40–46^. In contrast, none of the low-ranked HB genes has established roles in regulating influenza virus infection outcomes.

For high-ranked HB genes, we identified five unique virus-induced transcript expression clusters, and within each cluster, we observed HB genes with related biological functions (see **Fig. 1c**):

- **Cluster 1** (C1, light blue; upregulated by both pH1N1 and H5N1 infections) includes genes that are transcriptionally activated by NFκB (*TNF*, *TNFAIP3*, *CD69*, *SAA2*)^47^, suggesting roles in inflammatory responses. A portion of cluster members are regulators of inflammatory signaling (three are immunosuppressive), but other genes are less clearly linked to inflammation, with roles in regulating protein translation or lipid metabolism (**Fig. 1c**).
- **Cluster 2** (C2, pink; upregulated by pH1N1 and downregulated by H5N1 infection) includes multiple ISGs that are transcriptionally activated by pH1N1 but suppressed by H5N1 (*DDX58*, *IL4I1*, *ISG20*, *NMI*, *PARP12*, *TAP1*, and *ZNFX1*)^48, 49^. Cluster members have various roles in innate immune signaling in influenza virus infections and other contexts and include several potent influenza virus restriction factors and pathogenicity regulators (**Fig. 1c**).
- **Cluster 3** (C3, green; downregulated by both pH1N1 and H5N1 infections) includes genes that regulate cell fate, including cell cycle progression, cell differentiation, and apoptosis. Most cluster members lack known roles in influenza virus infection or innate immune signaling (**Fig. 1c**).
- **Cluster 4** (C4, orange; downregulated by pH1N1 and upregulated by H5N1 infection) includes three major regulators of membrane trafficking and secretion, of which two have clear pro-inflammatory functions. To date, none are linked to influenza virus infection or pathogenicity (**Fig. 1c**).
- **Cluster 5** (C5, gray; minimal expression changes in both pH1N1 and H5N1 infections) genes have no common functions, but include individual regulators of translation, DNA repair, or reactive oxygen species production (**Fig. 1c**).

In contrast, low-ranked HB genes exhibit less prominent expression changes in pH1N1 influenza virus infections and a distinct distribution among expression clusters (*i.e.*, few exhibit C1, C2, or C3 expression, while more than half are members of C5) (**Supplementary Fig. 1b**). These observations highlight substantial differences in the characteristics of high-ranked and low-ranked HB genes and support the notion that HB gene rankings partition host genes with unique regulatory effects on influenza virus infection and pathogenicity.

### An siRNA library for use in functional screening assays

To perturb the expression of HB genes, we used small interfering RNAs (siRNAs) to suppress the expression of HB genes (4 unique siRNAs per gene) (**Fig. 2a**). To ensure that siRNAs functioned correctly, we assayed siRNA effects on target gene knockdown and cellular viability and excluded siRNAs that did not appreciably reduce target mRNA expression (fold-change ≤ -1.4), or that caused a substantial reduction in intracellular ATP levels (≥ 30%), a proxy for loss of cellular viability. In all subsequent analyses, we included only HB genes for which ≥ 2 siRNAs targeting the same gene passed the target gene knockdown and viability criteria. In total, this comprised all 50 high-ranked HB genes and 9 (of 22) low-ranked HB genes (**Supplementary Table 2a**).

**Figure 2.**
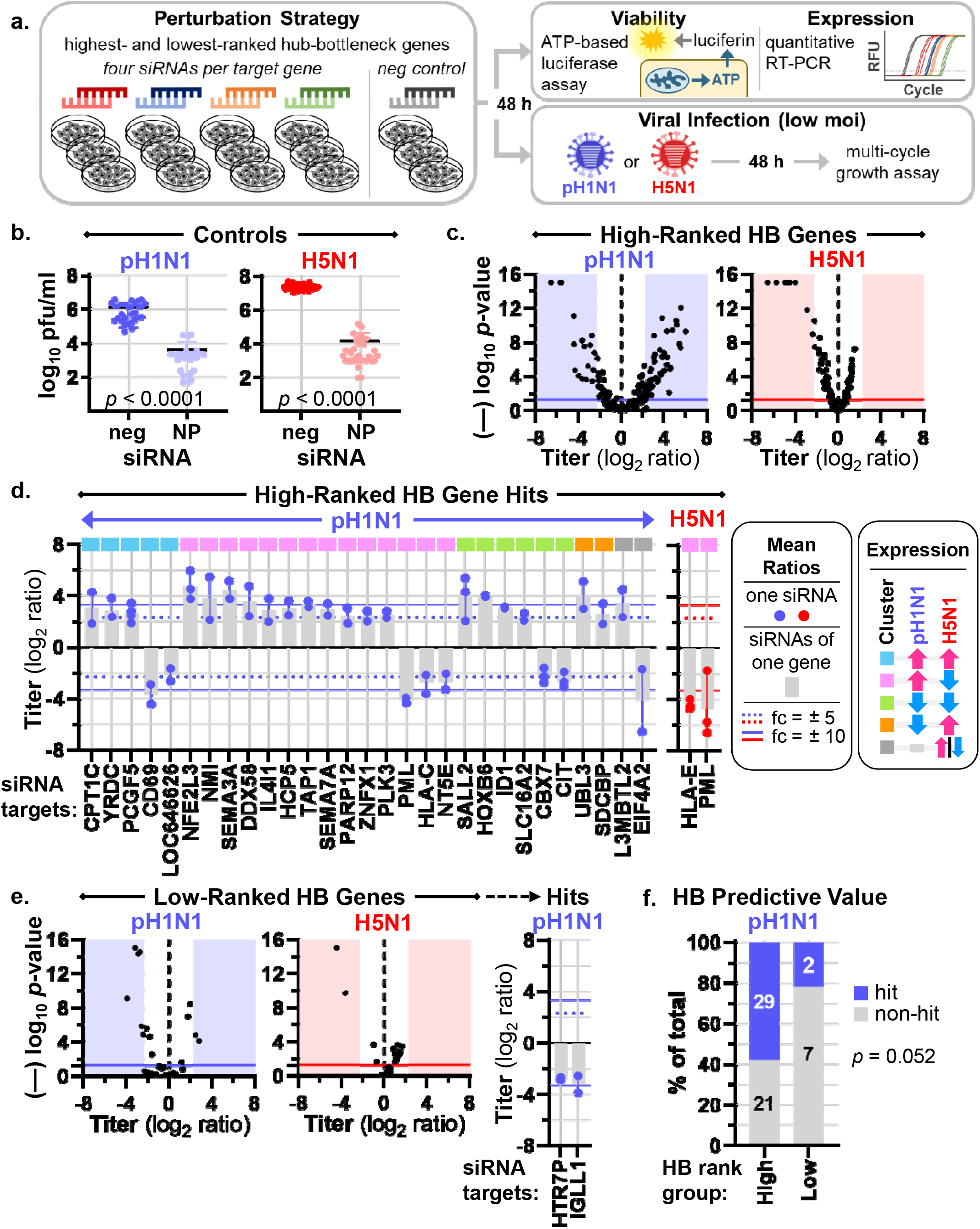
An RNAi screen to discover HB genes that regulate influenza virus growth. (**a**) The panel provides an overview of our siRNA-based strategy to perturb HB gene expression (**left**), and the phenotypic assays performed in siRNA-treated cells (**right**). (**b**) Graphs show pH1N1 (A/Oklahoma/VIR09-1170038L3/2009) (**left**) or H5N1 (A/Vietnam/1203/2004) (**right**) virus titers in A549 cells treated with negative control siRNA (neg) or an siRNA targeting the influenza nucleoprotein (NP) mRNA. Mean titers of negative control and NP siRNA-treated cells were compared by using a two-tailed, unpaired t-test with Welch’s correction, and the resultant *p*-values are indicated on the graphs. The data in the left and right panels represent 9 and 7 independent experiments, respectively. (**c)** Volcano scatter plots depict log_2_-transformed mean virus titer ratios (*x*-axis) versus (—) log_10_ *p*-values (*y*-axis) for cells treated with siRNAs targeting 50 high-ranked HB genes (2-4 siRNAs per gene) and assayed (in triplicate) for effects on pH1N1 (**left**) or H5N1 (**right**) virus growth (**Supplementary Table 2a**). To determine ratios, we compared mean titer values for HB gene siRNA- and negative control siRNA-treated cells, and *p*-values were calculated by using by using two-tailed, unpaired t-tests. In each plot, a data point represents the outcome for a single HB gene siRNA, the shaded areas demarcate ± 5-fold change in virus titer, and a solid horizontal line indicates *p* = 0.05. Each siRNA was evaluated in one experiment for each virus, and complete datasets were collected in a series of 5 and 3 independent experiments for pH1N1 and H5N1, respectively, and pooled to generate the plots. (**d**) The graphs summarize hit genes in pH1N1 (**left**) and H5N1 (**right**) virus growth assays, identified as described in **Supplementary Table 2b** by using the data shown in panel (**c**). Blue or red data points represent mean titer ratios for each individual siRNA contributing to the hit phenotype (derived from 2-4 siRNAs per HB gene and 3 replicate values per siRNA), gray bars represent grand mean titer ratios, and vertical connectors show the siRNA effect range. Hits are grouped according to expression clusters described in Fig. 1c, with cluster membership indicated at the top of the graph and a key shown at the right. fc, fold change. (**e**) We evaluated siRNAs targeting 9 low-ranked HB genes in pH1N1 and H5N1 virus growth assays and identified hit genes exactly as described for high-ranked HB genes. Volcano scatter plots (representing pooled data from 4 experiments for each virus) and the hit gene summary graph are presented as described in panels (**c**) and (**d**). (**f**) The graph shows the distribution of hits and non-hits for high- and low-ranked HB genes in the pH1N1 virus growth assay. The distributions were compared by using a one-sided Fisher’s Exact test, and the resultant *p*-value is shown on the panel.

### A high proportion of high-ranked HB genes are regulators of influenza virus growth

Next, we used siRNA passing the criteria described above to evaluate the effects of HB genes on pH1N1 or H5N1 virus replication. Importantly, for both pH1N1 and H5N1 infections, we observed consistent peak virus titers in cells treated with a non-targeting negative control siRNA, and consistently reduced virus titers (≥ 100-fold) in cells treated with a positive control siRNA targeting the influenza nucleoprotein (NP) mRNA (**Fig. 2b**). To assess the global effect of siRNAs targeting HB genes, we plotted the average log_2_ fold-change in virus titer (relative to non-targeting siRNA-treated cells) versus significance. For high-ranked HB candidate genes, scatter plots show a wide range of differences for both pH1N1 (fold-change range = -95.6 to +62.4) and H5N1 (fold-change range = -96.2 to +2.9), although changes in pH1N1 titers exhibit greater overall spread and more even distribution across positive and negative values compared to changes in H5N1 titers (**Fig. 2c** and **Supplementary Table 2a**). These observations suggested that high-ranked HB genes possess both pro-viral and antiviral factors.

To identify pro-viral and antiviral factors, we classified HB gene ‘hits’ from our virus replication datasets by using the following criteria: (*i*) the majority of siRNAs targeting a single HB gene must induce a statistically significant (*p* < 0.05) change in mean virus titer in the same direction; (*ii*) at least one of those siRNAs must induce ≥ (±) 6-fold change in virus replication; and (*iii*) the remaining siRNAs must induce ≥ (±) 3-fold change in virus replication (also see **Supplementary Table 2b**). Strikingly, we identified 29 high-ranked HB genes (58% of those evaluated) whose knockdown potently and consistently affected pH1N1 replication, including 21 anti-viral and 8 pro-viral hits (**Fig. 2d**, left panel). Among the hits are top-ranked HB genes previously shown to have anti-viral (*DDX58*)^8^ or pro-viral (*CIT*)^44, 50^ effects on influenza virus replication, and siRNAs targeting these genes resulted in the same outcomes in our assay, providing internal validation controls. Most of the other pH1N1 hits have no established link to influenza virus replication (**Fig. 1c**), strongly suggesting that HB nodes derived from transcriptional association networks are rich in novel pro-viral and antiviral regulators of influenza virus replication. Notably, hits encompassed HB genes from all five expression clusters (**Fig 2d**, see the colored squares at the top of the panel and compare to **Fig. 1c**).

When we applied the same selection criteria (**Supplementary Table 2b**) to our H5N1 siRNA dataset, we found fewer hits (2 of the 50 evaluated high-ranked HB genes; **Fig. 2d**, right panel). Although the reduced number of hits could indicate differences in host regulation of pH1N1 and H5N1 infection, it is also possible that the effects of siRNA treatment were obscured by higher replication levels in H5N1 infections (**Fig. 2b**) or are due to differences in virus-induced changes in HB gene expression (**Fig. 1c**). Therefore, for a subset of the high-ranked HB genes that exhibited antiviral effects in pH1N1 siRNA assays, we determined the effect of transient, cDNA-mediated ectopic expression on H5N1 multi-cycle replication. For three HB genes (*HOXB6*, *PCGF5*, and *SALL2*) out the six we tested, ectopic expression did not affect cell viability (**Supplementary Fig. 2a and 2b**), but significantly reduced H5N1 titers at either 24 h (**Supplementary Fig. 2c,** top) or 48 h (**Supplementary Fig. 2c,** bottom) post-infection. Therefore, high-ranked HB hits include novel regulators of both pH1N1 and H5N1 growth.

In contrast to high-ranked HB genes, few low-ranked HB genes affected pH1N1 or H5N1 replication; we identified only two genes with pro-viral effects in pH1N1 infections and none with antiviral effects (**Fig. 2e** and **Supplementary Table 2a**). Importantly, the hit-rate for high-ranked HB genes (58%) is higher than the hit rate for low-ranked HB genes (22.2%) (*p* = 0.052, Fisher’s exact test), indicating that high-ranked HB genes are enriched for influenza virus growth regulators (**Fig. 2f**). Altogether, our results indicate that HB rank-based predictions are predictive of host genes that regulate influenza virus replication and reveal multiple host genes with previously undiscovered pro-viral or antiviral effects.

### A subset of antiviral HB genes enhances cell autonomous innate immune signaling

To clarify how top-ranked HB genes exert their effects on influenza virus growth, we performed two additional, complementary siRNA screens to determine the effects of HB gene knockdown on regulation of cell autonomous innate immune signaling. For these studies, we developed A549 cell lines that stably express a luciferase reporter gene under the control of either the human *IFNB1* (A549-IFNB1-Luc) (**Supplementary Fig. 3a**) or NFκB-dependent (A549-NFκB-Luc) (**Supplementary Fig. 3b**) promoter. Importantly, induction of reporter gene expression in both cell lines is dependent on essential regulators of each cognate signaling pathway: DDX58, IRF3, and IRF7 for A549-IFNB1-Luc cells (**Supplementary Fig. 3a)**; or NFKB1 and RELA for A549-NFκB-Luc cells (**Supplementary Fig. 3b**).

To determine the effects of HB genes on *IFNB1* or NFκB-dependent gene expression, we transfected A549-IFNB1-Luc or A549-NFκB-Luc cells with siRNAs targeting HB genes and measured luciferase activity in mock (phosphate-buffered saline, PBS)-treated or stimulated cells (**Supplementary Table 3a** and **3b**). In A549-IFNB1-Luc cells treated with PBS or stimulated with polyinosine-polycytidylic acid (poly(I:C); a synthetic analog of dsRNA that activates antiviral PRRs [TLR3, DDX58, and EIF2AK2]), a positive control siRNA (targeting *IRF3* expression) consistently reduced luciferase activity compared to cells treated with the non-targeting negative control siRNA (*p* < 0.01) **(Supplementary Fig. 3c**, left panel [PBS-stimulated cells]; **Fig. 3a**, left panel [poly(I:C)-stimulated cells]). We observed a similar outcome in A549-NFκB-Luc cells treated with PBS or stimulated with TNF (which strongly activates NFκB) and treated with a positive control siRNA (targeting *RELA* expression) versus the non-targeting control (*p* < 0.001) (**Supplementary Fig. 3c**, right panel [PBS-stimulated cells]; **Fig. 3a**, right panel [TNF-stimulated cells]). To assess the global effects of siRNAs targeting HB candidates, we plotted the average log_2_ fold-change in luciferase activity (relative to non-targeting siRNA-treated cells) versus significance. For cells treated with siRNAs targeting HB genes and stimulated with PBS, this revealed minimal overall activation, demonstrating that siRNA treatment did not broadly stimulate cell autonomous innate immune signaling in a non-specific manner (**Supplementary Fig. 3d**). However, like the global effects of HB gene siRNAs on influenza virus replication (**Fig. 2c**), we observed a range of changes in luciferase activity in poly(I:C)-treated A549-IFNB1-Luc (fold-change range = -15.3 to +6.4) or TNF-treated A549-NFκB-Luc (fold-change range = -24.6 to +32.1) cells (**Fig. 3b**). We identified both positive and negative regulators among the hit genes for each pathway (see **Supplementary Table 3a** for A549-IFNB1-Luc data, **Supplementary Table 3b** for A549-NFκB-Luc data, and **Supplementary Table 3c** for reporter cell line hit selection criteria).

**Figure 3.**
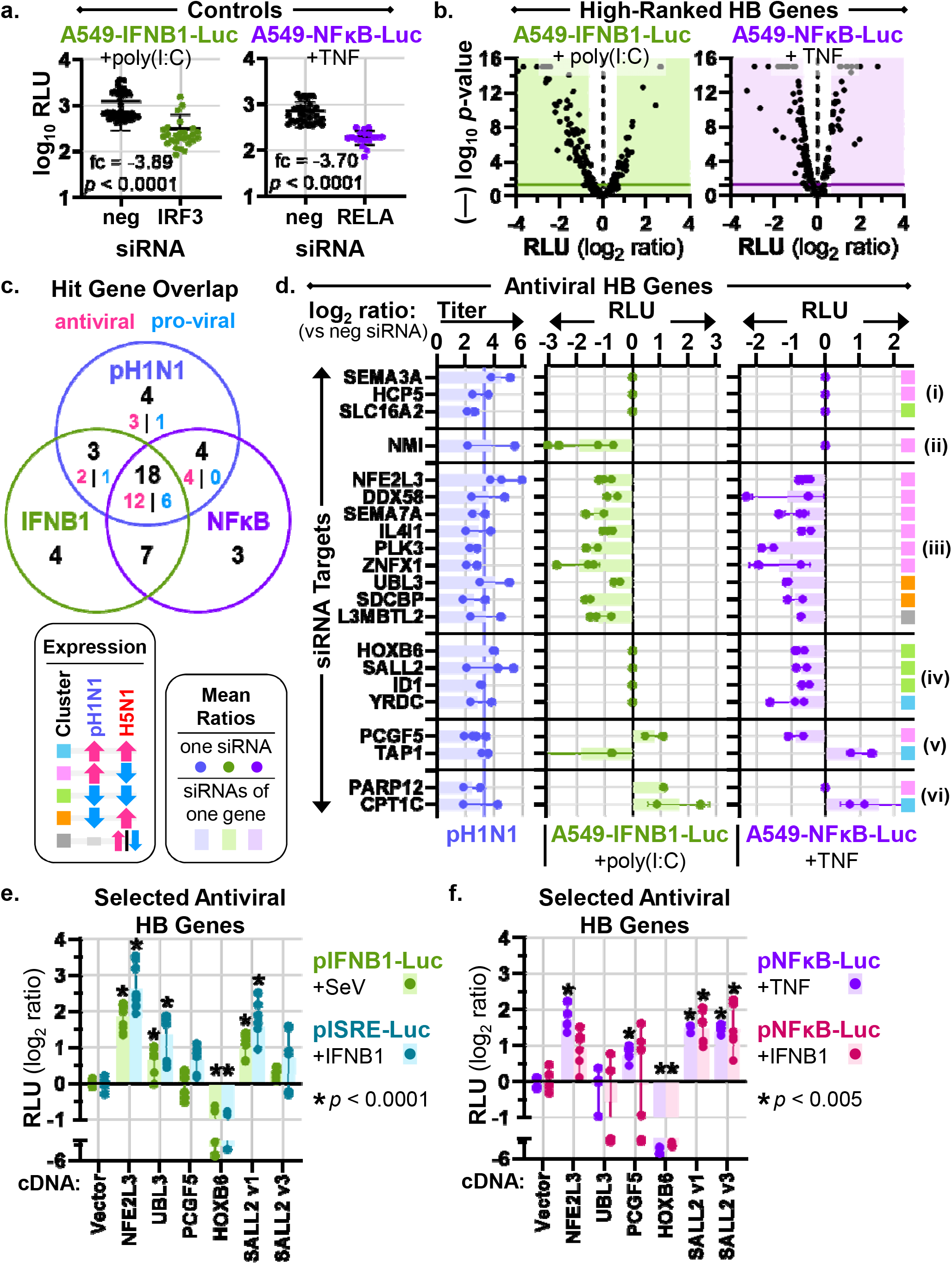
Complementary RNAi screens to identify HB genes that regulate cell autonomous innate immune signaling. We used siRNAs to perturb HB gene expression in luciferase reporter cell lines and then measured luciferase activity (relative light units, RLU) after stimulation. We used poly(I:C) to stimulate A549-IFNB1-Luc cells and TNF to stimulate A549-NFκB-Luc cells. (**a**) Graphs show RLU values in A549-IFNB1-Luc (**left**) or A549-NFκB-Luc (**right**) cells treated with control siRNAs and stimulated as described. Mean RLU values of cells treated with negative control siRNA (neg) and siRNA targeting expression of either *IRF3* or the *RELA* component of the NFκB transcription factor were compared by using two-tailed, unpaired t-tests, and the resultant fold changes and *p*-values are indicated on the graphs. The data in the left and right panels each represent 4 independent experiments. (**b**) Volcano scatter plots depict log_2_-transformed mean RLU ratios (*x*-axis) versus (—) log_10_ *p*-values (*y*-axis) for reporter cells treated with siRNAs targeting 50 high-ranked HB genes (2-4 siRNAs per gene), stimulated as indicated, and assayed (in triplicate) for effects on IFNB1 (**left**) or NFκB (**right**) activation (**Supplementary Table 3a** and **3b**). To determine ratios, we compared mean RLU values for HB gene siRNA- and negative control siRNA-treated cells, and *p*-values were calculated by using by using two-tailed, unpaired t-tests. In each plot, a data point represents the outcome for a single HB gene siRNA, the shaded areas demarcate ± 1.6-fold change in RLU, and a solid horizontal line indicates *p* = 0.05. Each siRNA was evaluated in one experiment in each reporter cell line, and complete datasets were collected in a series of 4 independent experiments for each cell line and pooled to generate the plots. (**c**) Hit genes in the IFNB1 and NFκB assays were identified as described in **Supplementary Table 3c** by using the data shown in panel (**b**). The Venn diagram shows the number of high-ranked HB gene hits that overlap between pH1N1, IFNB1, and NFκB assays (black text); and for pH1N1 hits, we designate the number of antiviral and pro-viral genes in pink and blue text, respectively (also see Fig. 2d). (**d**) The graph summarizes IFNB1 (green, **center**) and NFκB (purple, **right**) hits and non-hits for high-ranked HB genes exhibiting antiviral activity in the pH1N1 assay (21 genes; for ease of reference, mean titer ratios from the pH1N1 assay [see Fig. 2d] are shown in blue at the **left**). Blue, green, and purple data points represent mean ratios for each individual siRNA contributing to the hit phenotype (derived from 2-4 siRNAs per HB gene and 3 replicate values per siRNA), bars represent grand mean ratios, and vertical connectors show the siRNA effect range. A ratio value of 0 indicates a non-hit in either the IFNB1 or NFκB assays. Groups i—vi, (designated at the right) are discussed in the text of the **Results**. (**e**) & (**f**) Graphs show pooled RLU ratios of 293T cells ectopically expressing selected antiviral HB gene cDNAs along with pIFNB1-Luc or pISRE-Luc (3 independent experiments, 3 replicates per experiment) (**e**), or pNFκB-Luc (2 independent experiments, 3 replicates per experiment) (**f**). Cells were stimulated as indicated on the figure panels. Bars represent the grand mean RLU ratios for all replicate values of each HB gene cDNA, with individual replicates shown by circular data points and the effect range for each hit gene indicated by the vertical connectors. Mean RLU values of the negative control (“Vector”) and HB gene cDNA-treated cells were compared by using ordinary one-way ANOVA with a multiplicity adjustment, and significant

Next, we examined the overlap of hit genes identified from our pH1N1 replication, IFNB1 signaling, and NFκB signaling assays (**Fig. 3c**) and identified HB genes with signaling effects that might explain antiviral activity. Among high-ranked HB genes with antiviral activity, we observed six groups with unique IFNB1 and NFκB regulatory activity, of which a subset exhibited consistent effects on virus growth and cell autonomous innate immune signaling (**Fig. 3d**). Group 1 (**Fig. 3d** **(i)**) includes 3 antiviral genes (*SEMA3A*, *HCP5*, and *SLC16A2*) that did not affect IFNB1 or NFκB signaling in our assays, suggesting they may directly interfere with an essential step in influenza infection or affect another signaling pathway to repress replication. Group 2 (**Fig. 3d** **(ii)**) consists of a single antiviral gene (*NMI*) that positively regulates *IFNB1* expression (*i.e.*, knockdown suppresses luciferase activity), and Group 3 (**Fig. 3d** **(iii)**) comprises 9 genes (*NFE2L3*, *DDX58*, *SEMA7A*, *IL4I1*, *PLK3*, *ZNFX1*, *UBL3*, *SDCBP*, and *L3MBTL2*) that positively regulate both *IFNB1* and NFκB-dependent gene expression. Group 2 and group 3 genes may impose antiviral effects by promoting PRR activation, which stimulates both anti-viral (*IFNB1*) and pro-inflammatory (NFκB-dependent) gene expression. Group 4 (**Fig. 3d** **(iv)**) consists of 4 antiviral genes (*HOXB6*, *SALL2*, *ID1*, and *YRDC*) that positively regulate NFκB-dependent, but not *IFNB1*, gene expression. Group 4 genes may mediate antiviral effects through pro-inflammatory signaling independent of PRR activation. Group 5 (**Fig. 3d** **(v)**) includes 2 antiviral genes (*PCGF5* and *TAP1*) that positively regulate either *IFNB1* or NFκB-dependent gene expression and negatively regulate the other pathway; and group 6 (**Fig. 3d** **(vi)**) includes 2 antiviral genes (*PARP12* and *CPT1C*) that negatively regulate one or both pathways. Altogether, the antiviral activity of 14 high-ranked HB genes (groups 2, 3, and 4; **Fig. 3d** **(ii)**, **(iii)**, and **(iv)**) may be explained by their ability to positively regulate antiviral and/or pro-inflammatory signaling pathways; with the exception of *DDX58*^8^ and *ZNFX1*^38^, none is reported to regulate innate immune signaling in influenza virus-infected cells. Regarding high-ranked HB genes with pro-viral activity (8 in total, **Fig. 2d** and **3c**), only 2 exhibit consistent effects on viral growth and cell autonomous innate immunity (*i.e.*, knockdown suppresses virus growth and upregulates luciferase activity in reporter cells): *CIT* negatively regulates IFNB1 signaling and *CD69* negatively regulates both IFNB1 and NFκB signaling (**Supplementary Table 3**). Therefore, CIT and CD69 may promote virus replication by suppressing PRR signaling. While CIT is reported to have pro-viral activity in influenza infections^44, 50^, it is not known to regulate innate immune signaling pathways.

To confirm the ability of high-ranked HB genes to regulate cell autonomous innate immune signaling, we transiently expressed cDNAs of a subset of antiviral HB genes (*NFE2L3* and *UBL3*, group 3; *HOXB6* and *SALL2*, group 4; and *PCGF5*, group 6)—along with plasmids encoding *IFNB1*, NFκB-dependent, or IFN-stimulated response element (ISRE) promoter-reporter cassettes—and measured luciferase activity in stimulated cells. We confirmed positive regulation of *IFNB1* expression for NFE2L3 and UBL3 and showed that one variant of SALL2 has similar regulatory effects (**Fig. 3e**). Ectopic expression of all three of these cDNAs also upregulated ISG expression (determined by ISRE-dependent luciferase reporter activity), the direct downstream target of type I IFN, further supporting their ability to positively regulate antiviral signaling (**Fig. 3e**). NFE2L3, PCGF5, and both SALL2 variants (but not UBL3) also positively regulated NFκB-dependent gene expression activated by either TNF or IFNB1 (**Fig. 3f**). Collectively, these analyses suggest that high-ranked HB genes regulate influenza virus replication by modulating the activation of host antiviral and inflammatory signaling pathways, which is consistent with their predicted ability to regulate the host transcriptional network in response to influenza virus infection.

### Hub-bottleneck genes exhibit partially overlapping regulation of diverse viruses

As many of the pro-viral and antiviral genes identified in our siRNA screens have no identified roles in regulating virus replication, we wondered whether they might also regulate infection of other viruses. Therefore, we selected 18 high-ranked HB gene hits with the following characteristics for additional studies with other viruses: they (***i***) represent HB gene transcriptional expression clusters C1—C4; (***ii***) have high antiviral or pro-viral activity in influenza virus infection; (***iii***) exhibit a variety of innate immunity regulation phenotypes; and (***iv***) include factors with no known role in regulating virus replication or innate immune signaling. Virus growth, IFNB1, and NFκB-dependent gene expression phenotypes of the selected HB gene hits are summarized in **Supplementary Table 4a**.

We assayed the effect of siRNA-mediated knockdown of the 18 selected HB genes on multi-cycle replication of four additional viruses, including an H3N2 influenza strain (A/Yokohama/2017/2003), a biologically-contained Ebola-ΔVP30 virus (EBOV)^51^, West Nile virus (WNV), and Middle East respiratory syndrome coronavirus (MERS-CoV) (**Supplementary Table 4b**, **4c**, and **4d**). This diverse group includes other respiratory pathogens (H3N2 and MERS-CoV), a neurotropic virus (WNV), and a systemic virus (EBOV); as well as viruses with either negative-sense (H3N2 and EBOV) or positive-sense (WNV and MERS-CoV) RNA genomes, each possessing unique strategies for genome replication. As expected, most high-ranked HB genes regulated H3N2 replication in a manner similar to pH1N1 (**Fig. 4a** and **4b**), suggesting multiple genes with conserved effects across different influenza viruses. Remarkably, the effects of more than half of the evaluated high-ranked HB genes were similar between pH1N1 or H3N2 and EBOV (**Fig. 4b**), suggesting that a subset of pro-viral and antiviral HB genes could similarly affect the replication of some negative-sense RNA viruses. We observed less similarity between influenza viruses and WNV, although several genes exhibited overlapping pro-viral activity for WNV, influenza viruses, and EBOV (*PML*, *HLA-C*, *HLA-E*, and *ARL3*) (**Fig. 4b**). There was little similarity in functional activities of high-ranked HB genes in MERS-CoV infection and any other virus tested here (**Fig. 4b**). Therefore, high-ranked HB genes identified from influenza virus infections have partially overlapping regulatory effects on growth of diverse viruses, suggesting some could be exploited for development of broad-spectrum antiviral treatments; and that in-depth analysis of host genes with unique effects on growth of different viruses may reveal important differences in viral pathogenicity mechanisms.

**Figure 4.**
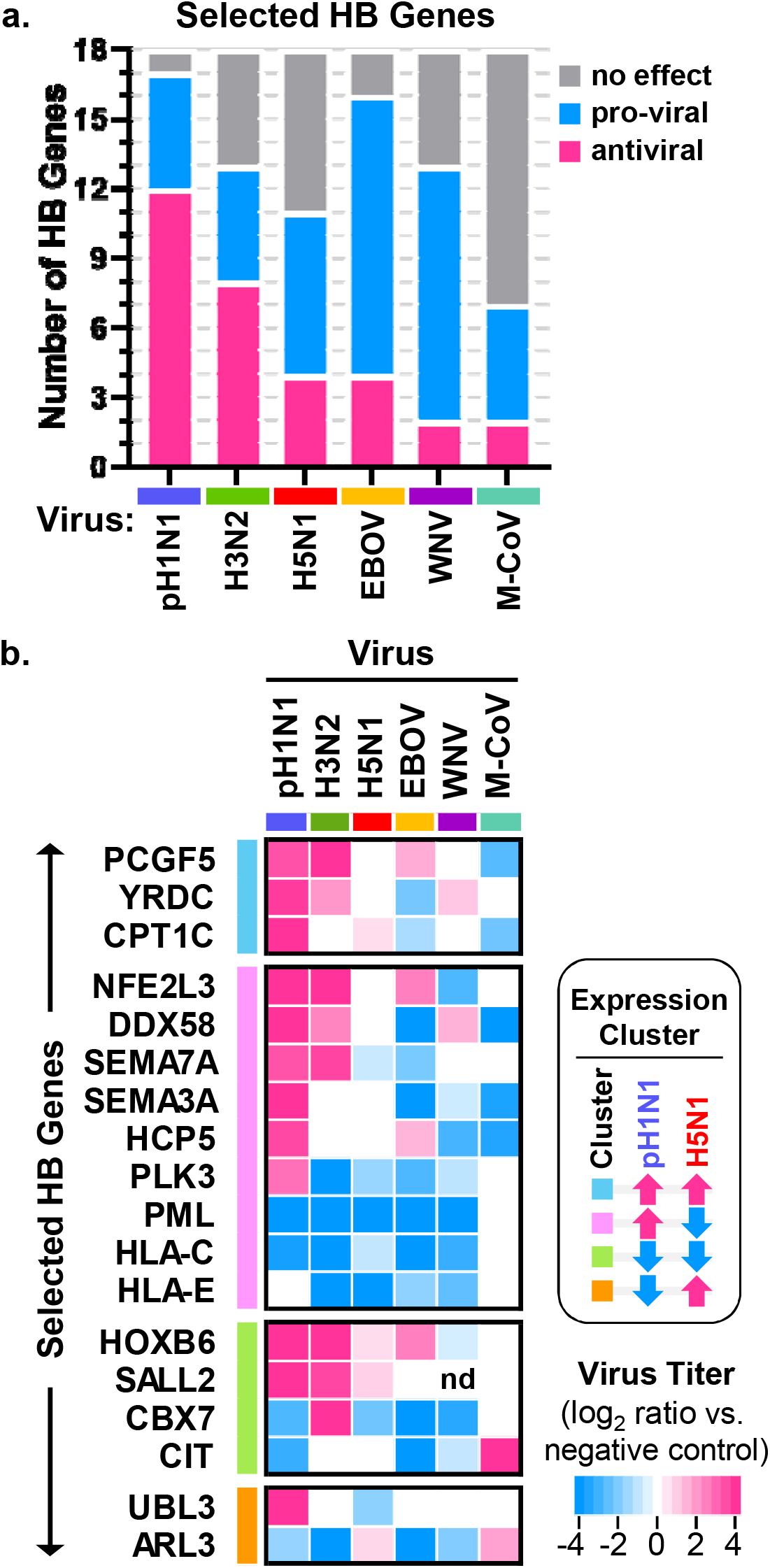
Effects of high-ranked HB genes on growth of diverse viruses. (**a**) For a selected set of 18 high-ranked HB genes that regulate pH1N1 virus growth, we determined the effects of siRNA treatment on growth of four other viruses: an H3N2 influenza virus (‘H3N2’), Ebola-ΔVP30 virus (‘EBOV’), West Nile virus (‘WNV’), and Middle East respiratory syndrome coronavirus (‘M-CoV’) (see **Supplementary Tables 4a** and **4b**). For each virus tested, the graph shows the number of hit genes with pro-viral or anti-viral effects and the number of non-hit genes with no effect on virus growth (hit selection criteria are described in **Supplementary Table 4c**). Corresponding numbers of pro-viral, antiviral, and non-hit genes from the pH1N1 and H5N1 siRNA screens (see **Supplementary Table 2a**) are given for comparison. (**b**) For each virus, the heat map shows the log_2_ mean virus titer ratio (versus negative control) for the active siRNA with the strongest effect on virus replication (a heat map key is given at the lower right). If none of the siRNAs targeting a single gene met the hit selection criteria (**Supplementary Table 4c**), then the log_2_ mean virus titer ratio is represented as 0 (white) on the heat map. For H3N2, EBOV, and M-CoV, each siRNA was evaluated in triplicate in one experiment, and complete datasets were collected in a series of independent experiments (H3N2 and EBOV, 2 independent experiments per virus; M-CoV, 5 independent experiments). For WNV, each siRNA was evaluated in two independent experiments with 3-4 replicates per experiment. Data from pH1N1 and H5N1 screens (see **Supplementary Table 2a**) are given for comparison. HB target genes are grouped according to the expression clusters identified in Fig. 1c, which are indicated by the colored bars at the left of the heat map and a key at the right.

### Nfe2l3 regulates pH1N1 replication and cytokine production *in vivo*

In our final set of experiments, we examined whether a high-ranked HB gene with effects on influenza virus growth and host response signaling *in vitro* similarly regulated these phenotypes *in vivo*. We focused on NFE2L3, which exhibited potent antiviral effects (**Fig. 2b**), positive regulation of type I IFN and NFκB activation at levels similar to that of DDX58 (**Fig. 3d-f**), and conserved antiviral activity against two influenza virus strains and EBOV *in vitro* (**Fig. 4b**). We inoculated female wild-type (WT) and *Nfe2l3* knockout (KO)^52^ mice with a sub-lethal dose of pH1N1 (10^4^ plaque-forming units [pfu] per mouse) and monitored body weight for 16 days. Whereas WT and KO mice exhibited similar body weight loss over the first 8 days after infection, average body weights diverged during the latter 8 days of infection as WT mice recovered more quickly (**Fig. 5a**). A mixed effects model indicated that average daily body weights were significantly different between WT and KO mice on days 13-16 post infection (*p* < 0.05). These results are consistent with the possibility that Nfe2l3 has protective antiviral effects *in vivo*.

**Figure 5.**
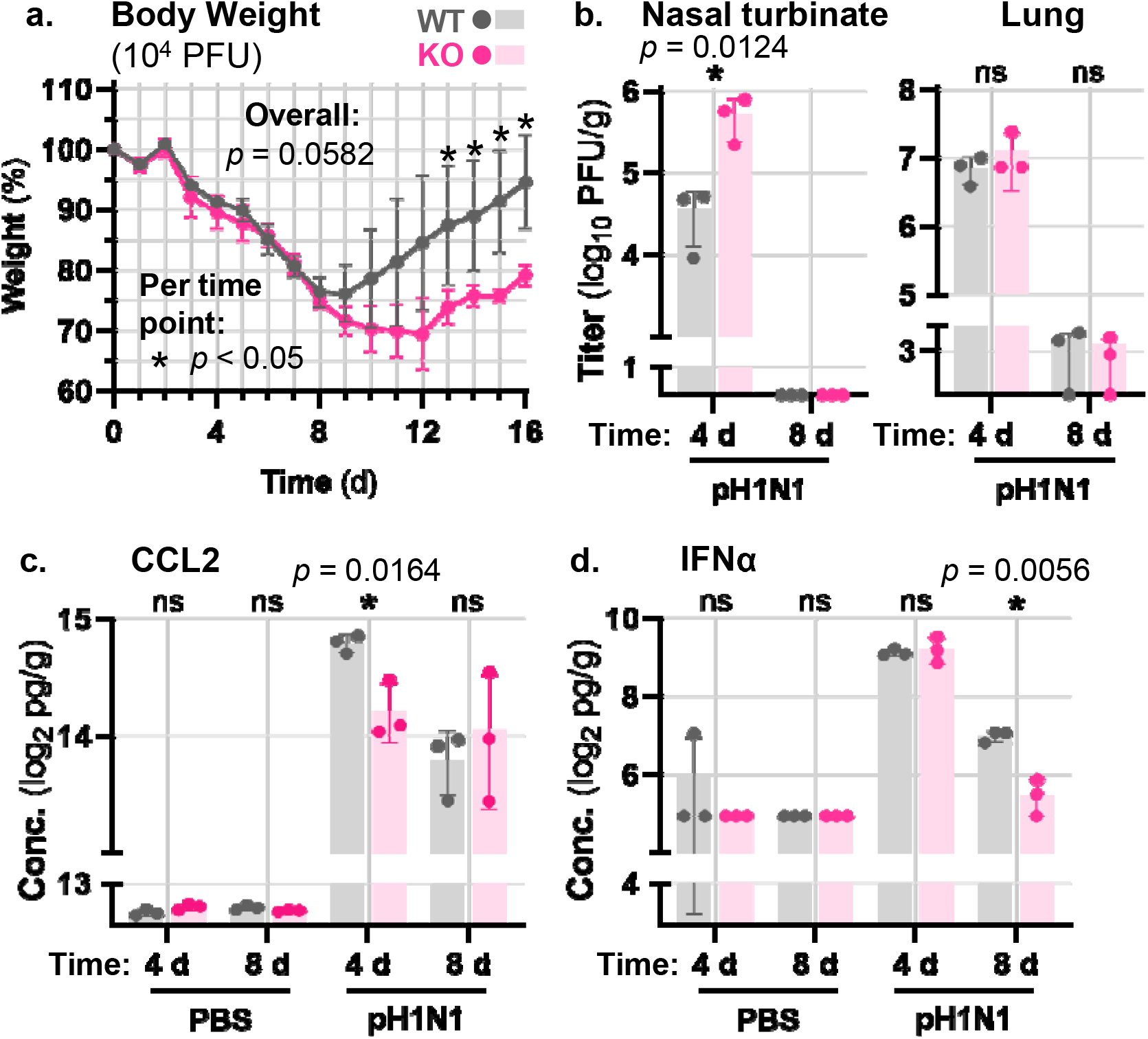
Influenza pathogenesis in *Nfe2l3* knockout mice. (**a**) Groups of female wild-type (WT; n = 4) or *Nfe2l3* knockout (KO; n = 3) mice were inoculated with 10^4^ pfu of influenza A/Oklahoma/VIR09-1170038L3/2009 (H1N1) and body weights were measured daily for 16 days. The graph depicts the average body weights for each group (expressed as a percentage of the body weight at the start of the experiment), with variation represented by standard deviation. Group weight loss profiles were compared by using a mixed effects model with a Geisser-Greenhouse correction, which generated a *p*-value for the overall effects of *Nfe2l3* expression on mouse body weight loss, as well as *p*-values for effects observed at each time point (both are given on the figure panel). One *Nfe2l3* KO mouse succumbed to the infection at 13 days post-inoculation. (**b**) **—**(**d**) Groups of female WT or *Nfe2l3* KO mice were inoculated with phosphate-buffered saline (PBS) or 10^4^ pfu of influenza A/Oklahoma/VIR09-1170038L3/2009 (H1N1) and euthanized on day 4 or day 8 post-inoculation for respiratory tissue collection (3 mice were inoculated with PBS or influenza virus for each mouse strain at each time point). Panel (**b**) shows virus titers in nasal turbinate and lung tissues in pH1N1 infections; and panel (**c**) and (**d**) show CCL2 or IFNα expression, respectively, in lungs of PBS-treated or pH1N1-infected mice. Error bars represent standard deviation, and *p*-values were calculated by using unpaired, two-tailed t-tests, and significant values (*p* < 0.05) are indicated on the figure panels. ns, not significant. The data in panel (**a**) and panels (**b**)—(**d**) were collected in 2 independent experiments.

To determine whether Nfe2l3 regulates virus growth and activation of antiviral and pro-inflammatory signaling pathways *in vivo*, we inoculated female WT and KO mice with pH1N1 (10^4^ pfu) and collected tissues at 4 and 8 days post-infection for virus titration (nasal turbinates and lungs) and multiplex cytokine analysis (lungs). Similar to the effects of NFE2L3 *in vitro*, we observed increased pH1N1 virus titers in nasal turbinate tissues at day 4 post-infection in mice lacking *Nfe2l3* expression (**Fig. 5b**; *p* = 0.0124), and reduced CCL2 (a pro-inflammatory cytokine expressed by airway epithelial cells in influenza virus infection^6, 7^) and type I IFN in lung tissues at days 4 and 8 post-infection, respectively (**Fig. 5c** and **5d**; *p* < 0.05). Together, these results indicate that Nfe2l3 possesses antiviral activity and promotes antiviral and inflammatory gene production *in vivo* and suggest that antiviral effects are mediated through cell autonomous innate immune signaling networks that activate type I IFN and NFκB-dependent gene expression. Therefore, we have demonstrated that a candidate HB regulator predicted by computational analysis of host transcriptional network responses *in vitro* promotes similar phenotypes and regulates influenza virus pathogenicity *in vivo*.

## DISCUSSION

Influenza virus disease severity is driven by virus growth in target tissues and host-dependent immunopathology. To identify key host factors that modulate these pathophysiological processes, we systematically tested computationally predicted regulators of influenza infection outcomes in experimental systems. In our prior work, we identified and prioritized candidate host regulators by using the topological features of a transcriptional association network model (*i.e.*, genes with both high hub-like and high bottleneck-like activity, or HB genes)^21^. Here, we validated dozens of high-ranked HB candidate genes (predicted by our model) as *bona fide* regulators of influenza virus growth and demonstrated that most HB genes mediate their effects through activation of IFNB1 and/or NFκB-dependent gene expression. We extended these findings by determining the activity of selected antiviral and pro-viral HB genes in infections with a set of unrelated RNA viruses, which indicated that some HB genes have partially overlapping growth regulatory activity across multiple virus families. We also demonstrated that a high-ranked antiviral HB gene (*NFE2L3/Nfe2l3*) similarly regulated virus replication, IFNB1 activation, and NFκB-dependent gene expression in human cells and mouse lung, and that this HB gene limits influenza disease severity in mice. Collectively, our results validate our HB node-based candidate regulator prioritization strategy, underscore the biological importance of high-ranked HB genes in influenza virus pathogenicity, and identify host factors with previously unknown roles in regulating virus growth and cell autonomous innate immunity.

A principal goal of this study was to test the hypothesis that high-ranked HB genes are more likely to possess key roles in influenza infection outcomes compared to low-ranked HB genes, and our findings unambiguously support this concept. High-ranked HB genes comprise a substantial group of host factors that prominently and consistently regulate influenza virus growth, including 21 antiviral and 8 pro-viral genes (29 hits out of 50 tested genes, or a hit rate of 58%), and are enriched for influenza virus growth regulators compared to low-ranked HB genes. Among the 29 high-ranked HB genes with antiviral or pro-viral activity, 16 also regulate cell autonomous innate immune signaling activity in a manner consistent with their effects on virus replication (14 antiviral HB genes and 2 pro-viral HB genes), and most have no previously reported role in regulating virus growth or virus-induced antiviral or inflammatory responses. Thus, our findings establish that prioritizing host genes by HB node rankings is an effective strategy for identifying regulators of influenza infection outcomes, and we suggest that similar strategies could be implemented to further explore influenza disease mechanisms or to investigate other infectious or non-infectious diseases where immunopathology has a central role in pathogenesis.

While a few of the novel regulators of influenza virus growth and cell autonomous innate immune signaling have been linked to regulation of antiviral or inflammatory signaling in other contexts (NFE2L3, NMI, SEMA7A, SDCBP)^52–60^ or adaptive immunity (IL4I1, ID1, CD69, UBL3, PLK3)^61–65^, others have no links to immune responses (L3MBTL2, YRDC, HOXB6, SALL2, and CIT). Determining how these host factors regulate antiviral and pro-inflammatory signaling to control influenza virus growth may lead to insights into influenza disease pathogenesis, as well as a better understanding of the pathways and factors that regulate host innate immunity. For example, several antiviral HB genes that activate both IFNB1 and NFκB-dependent gene expression (likely via PRR signaling, *e.g.*, RIG-I), also regulate DNA damage responses (*i.e.*, NFE2L3^66^, L3MBTL2^67^, and PLK3^68^). Recent studies have revealed links between DNA damage and innate immunity^69–72^, including a crucial role for the non-homologous end-joining protein, XRCC4, in potentiating type I IFN expression via RIG-I and protecting against influenza pathogenicity in the mouse model^73^. We suggest that NFE2L3, L3MBTL2, and/or PLK3 may represent additional links between DNA damage and innate immune signaling in response to viral infection, a possibility that needs to be examined more thoroughly.

Antiviral HB genes that have different effects on innate immune signaling also have different functional attributes (summarized in **Fig. 6**). For example, HB genes that induce antiviral activity via PRR signaling (positively regulating both IRF3/IRF7 and NFκB) (**Fig. 6**, center) are members of HB gene expression clusters C2 (pink), C4 (orange), or C5 (gray), which are differentially expressed between pH1N1 and H5N1 infections (*i.e.*, upregulated in one condition and downregulated [or not induced] in the other); are widely localized among different cellular organelles (including the nucleus, cytoplasm, Golgi apparatus, mitochondria, plasma membrane, and extracellular exosomes); and possess diverse biological functions. In contrast, HB genes that may promote antiviral activity by activating pro-inflammatory signaling independent of PRRs (positively regulating only NFκB and not IFNB1) (**Fig. 6**, left) are members of HB expression clusters C1 (light blue) or C3 (green), which are similarly expressed in both pH1N1 and H5N1 infections (*i.e.*, upregulated or downregulated in both conditions), and comprise mostly nuclear factors with established roles in regulating non-immune gene expression. Although the implications of these observations are presently unclear, we suggest that virus-induced expression patterns, cellular localizations, biological functions, and/or phenotypic effects of high-ranked HB genes may be linked, and that this phenomenon might be exploited to improve network-based predictions of candidate regulators in future studies.

**Figure 6.**
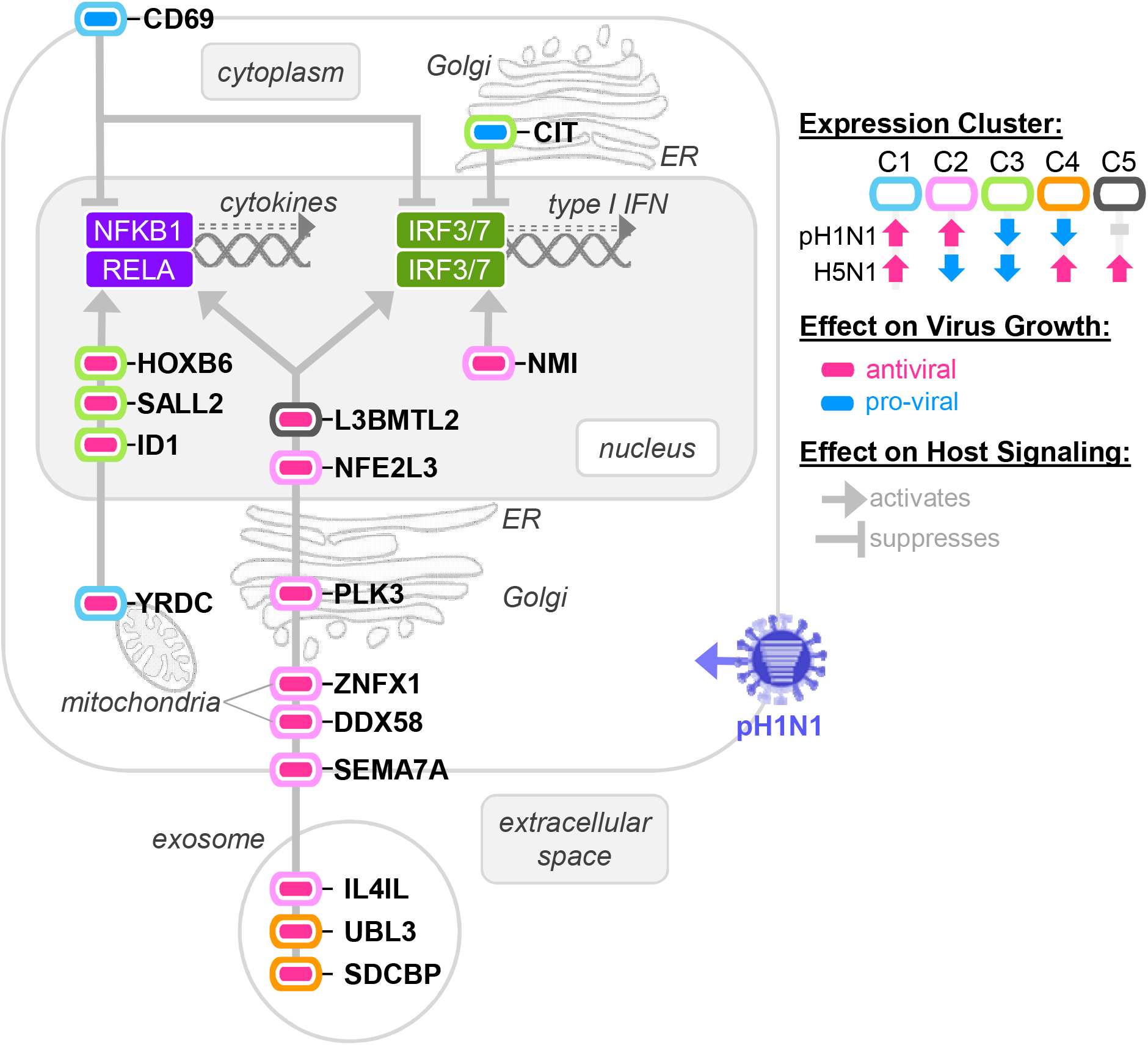
Functional summary of high-ranked HB genes that consistently regulate influenza virus growth and cell autonomous innate immune signaling. For high-ranked HB genes with pro-viral or antiviral activity that can be explained by effects on IFNB1 and/or NFκB activation we summarize expression in influenza virus-infected cells, cellular localization, effect on virus replication, and effects on antiviral and pro-inflammatory gene expression. HB genes centered on a single vertical gray bar have similar effects on IFNB1 and NFκB signaling. A key is provided at the right. Some HB gene proteins may exhibit additional subcellular localizations not shown in the figure. ER, endoplasmic reticulum.

In summary, we have demonstrated that transcriptional association network topology predicts host factor control of influenza virus growth through regulation of cell autonomous innate immune signaling, and we have identified a high priority list of virus growth and innate immunity regulators for in-depth analysis in future studies. Ultimately, this work expands our knowledge of the relationships between host transcriptional networks and regulation of influenza virus growth and virus-induced disease and may facilitate future development of antiviral countermeasures.

## MATERIALS AND METHODS

All assays associated with commercially available kits or reagents were carried out according to the manufacturers’ instructions.

### Bioinformatics

We employed the web-based Metascape portal (http://metascape.org)^27^ to infer enriched GO biological processes, KEGG pathways, Reactome gene sets, and canonical pathways in high-ranked and low-ranked HB gene sets. To assign individual functions of HB genes, we manually integrated Metascape results and information obtained by literature mining in the Entrez Pubmed database (https://pubmed.ncbi.nlm.nih.gov/).

### Microarray datasets

Transcript expression data are derived from the same datasets that we previously used for HB gene identification and were statistically processed previously^21, 48, 74^. Briefly, genome-wide changes in transcript expression were determined by microarray analysis (Agilent Technologies, Inc.) in human bronchial epithelial cells (Calu-3) infected with pH1N1 (A/California/04/2009) (multiplicity of infection [moi] = 3 plaque-forming units [pfu] per cell; 0, 3, 7, 12, 18, 24, 30, 36, and 48 h post-infection) or H5N1 (A/Vietnam/1203/2004) (moi = 1 pfu per cell; 0, 3, 7, 12, 18, 24 h post-infection), and log_2_ expression ratios and false-discovery (FDR) rate-adjusted q-values were determined compared to time-matched mock-infected controls. The log_2_ expression ratios and FDR-adjusted q-values of the selected high- and low-ranked HB genes are provided in **Supplementary Table 1e** and **Supplementary Table 1f**, respectively.

### Cell lines

We purchased A549 (human lung epithelial), 293T (human embryonic kidney epithelial), Huh7 (human hepatocellular carcinoma), BHK-21 (Syrian golden hamster kidney fibroblast), and Vero (African green monkey kidney epithelial) cell lines from the American Type Culture Collection (ATCC), and we used an in-house clone of the Madin-Darby canine kidney (MDCK) cell line adapted to grow in newborn calf serum (NCS) and support efficient influenza virus replication. To propagate cell lines, we used Eagle’s minimum essential medium (MEM) containing 5% NCS (MDCK); a 1:1 mix of Dulbecco’s modified Eagle’s medium (DMEM) and Ham’s F12 medium (DMEM/F12) containing 10% fetal bovine serum (FBS) (A549); DMEM containing 10% FBS (293T and Huh7); or MEM containing 10% FBS (BHK-21 and Vero). We also used Vero 76 and Huh7 cells stably expressing the Ebola VP30 protein (Vero-VP30 and Huh7-VP30 cells, respectively), described previously^51, 75^, which we propagated in MEM containing 10% FBS (Vero-VP30) or DMEM containing 10% FBS (Huh7-VP30). We supplemented all cell propagation media with L-glutamine and antibiotics, and cell lines were grown at 37°C in an atmosphere of 5% CO_2_. To ensure experimental reproducibility, we routinely monitored cell cultures for mycoplasma contamination, and periodically restarted cell lines from cryopreserved early passage aliquots.

### Viruses

The pH1N1 and H5N1 virus strains used throughout the study included influenza A/Oklahoma/VIR09-1170038L3/2009 (H1N1; ‘OK38L3’) and influenza A/Vietnam/1203/2004 (H5N1; ‘VN1203’), originally provided by the United States (US) Centers for Disease Control and Prevention (CDC)). We generated pH1N1 and H5N1 virus stocks by passaging the original isolate (OK38L3) or a reverse genetics supernatant (VN1203)^76, 77^ once in MDCK cells, as previously described^78^, and the same virus stocks were used for all siRNA screening, cDNA ectopic expression, or mouse infection experiments. In some siRNA experiments, we used the following additional viruses: a seasonal H3N2 influenza virus strain (A/Yokohama/2017/2003; ‘YK2017’)^79^; a biologically contained EBOV (Ebola-ΔVP30), which expresses green fluorescent protein in the place of the essential Ebola VP30 protein, and replicates only in cells expressing the Ebola VP30 protein^51^; a recombinant molecular clone of MERS-CoV based on the EMC2012 strain^80^; and WNV New York 1999 clone 382-99^81^. We generated virus stocks by passaging reverse genetics supernatants (H3N2, Ebola-ΔVP30, MERS-CoV), as previously described^51, 78–80^; or by electroporation of *in vitro* transcribed RNA into BHK-21 cells, as previously described^82, 83^, followed by passaging once in Vero cells (WNV). In other studies, we generated a Sendai virus stock (Enders strain) by passaging the isolate once in 10-day-old embryonated chicken eggs. To quantify all stock virus titers, we used standard plaque assays or focus forming unit (ffu) assays, as appropriate.

### siRNAs

We included the same non-targeting negative control (AllStars Negative Control siRNA, Qiagen) and transfection efficiency control (AllStars Hs Cell Death Control siRNA, Qiagen) in all siRNA assays. The non-targeting control has no homology to any known mammalian gene, elicits minimal non-specific effects, and does not affect influenza virus replication. The transfection efficiency control is blend of highly potent siRNAs targeting ubiquitously expressed human genes that are essential for cell survival and induces a cell death phenotype visible by light microscropy within 48 h (293T and A549 cells) or 72 h (Huh7 cells) (in all three cell types, this phenotype manifests as rounded cell morphology and detachment from the monolayer). We also used the following positive control siRNAs in appropriate assays: a previously described siRNA (NP-1496; referred to as ‘NP’ in the figures) that targets influenza nucleoprotein mRNA and inhibits influenza virus replication^84, 85^ (we provided the sequence to Integrated DNA Technologies [IDT], Inc., for synthesis); a pre-designed siRNA targeting the cellular *NPC1* gene, which inhibits EBOV replication (NPC1 is required for efficient Ebola virus entry^86, 87^); pre-designed siRNAs targeting cellular genes required for type I IFN activation (*DDX58*, *IRF3*, and *IRF7*); and pre-designed siRNAs targeting cellular genes required for NFκB activation (*RELA* and *NFKB1*). For the 50 high-ranked HB genes and 22 low-ranked HB genes, we obtained FlexiTube GeneSolution siRNA sets (Qiagen), comprising four unique, pre-designed siRNAs targeting each candidate gene mRNA (288 individual siRNAs in total). A list of all siRNAs (including product numbers and sequences, where applicable and available) is provided in **Supplementary Table 5a**.

### Plasmids

For a subset of high-ranked HB hit genes, we performed complementary gene perturbation studies in cells transiently overexpressing exogenous HB gene cDNAs. We purchased plasmids expressing these cDNAs, as well as a negative control plasmid (pCMV6-Entry) lacking a cDNA insert (OriGene Technologies, Inc.). A list of all plasmids encoding HB gene cDNAs (including product numbers, vector backbones, epitope tag placements, and the predicted molecular weights of proteins derived from the cDNAs) are provided in **Supplementary Table 5b**. In other studies, we used plasmids encoding cellular promoter-driven firefly luciferase reporter cassettes to assay the activation of type I IFN expression (pIFNB1-Luc), ISG expression (pISRE-Luc), or NFκB-dependent gene expression (pNFκB-Luc). We purchased the pISRE-Luc and pNFκB-Luc plasmids (Stratagene Products Division, Agilent Technologies, Inc.). To create pIFNB1-Luc, we PCR-amplified the human *IFNB1* gene promoter (including 125 base pairs upstream of the ATG start site) from Huh7 genomic DNA and inserted the promoter element upstream of the firefly luciferase open reading frame in the pGL2-Basic vector (Promega Corp.). In all cases, we isolated fresh plasmid DNA for use in transfection experiments by using the PureLink^TM^ HiPure Plasmid Maxiprep Kit (Invitrogen, Thermo Fisher Scientific Corp.). This method minimizes lipopolysaccharide contamination in plasmid DNA extracted from *Escherichia coli* and reduces non-specific PRR pathway activation in plasmid DNA transfections of mammalian cells. In addition, we used Sanger sequencing to verify that plasmids encoded the correct cDNA inserts, in-frame epitope tags, and/or promoter-luciferase cassettes prior to use in any experiments.

### Reporter cell lines

For siRNA screening studies, we created A549 cell lines stably expressing *IFNB1* promoter-luciferase or NFκB promoter-luciferase cassettes (‘A549-IFNB1-Luc’ or ‘A549-NFκB-Luc’, respectively). Briefly, we co-transfected low-passage A549 cells with pIFNB1-Luc or pNFκB-Luc and a linear hygromycin DNA fragment flanked by the SV40 promoter and polyadenylation signal (Clontech Laboratories, Inc.) by using the TransIT LT-1 transfection reagent (Mirus Bio LLC). Two days later, we trypsinized the transfected cells, plated them under limiting dilution, and carried out selection by continuously culturing cells in 1:1 DMEM/F12 containing 10% FBS, glutamine, an antibiotic mixture, and 500 μg/ml hygromycin (Sigma-Aldrich). Subsequently, we screened hygromycin-selected cell clones for the appropriate pathway activation, and for each reporter cell type, identified a clone exhibiting ≥ 10-fold increased luciferase activity after stimulation, and cryo-preserved low-passage aliquots for future use. To propagate A549-IFNB1-Luc and A549-NFκB-Luc cells, we used 1:1 DMEM/F12 containing 10% FBS, glutamine, a penicillin/streptomycin mixture, and 200 μg/ml hygromycin.

### siRNA and plasmid DNA transfections

We performed transfections for all phenotypic assays in 24-well plates seeded with 8 x 10^4^ cells per well. For siRNAs, we transfected cells two hours after seeding by using the Lipofectamine RNAiMAX reagent (Invitrogen, Thermo Fisher Scientific Corp.) and 20 nM (final concentration per well) of a single siRNA. For plasmids, we transfected cells 16 h after seeding by using FuGene HD transfection reagent (Promega Corp.) and either 0.2 µg of DNA (for plasmids encoding HB gene cDNAs used in multi-cycle influenza virus growth assays) or 0.6 µg of DNA (0.5 µg of plasmids encoding HB gene cDNAs plus 0.1 µg of pIFNB1-Luc, pISRE-Luc, or pNFκB-Luc for luciferase reporter assays, viability assays, and immunoblotting). To allow for target gene knockdown or protein accumulation prior to initiating phenotypic analyses, we incubated cells with plasmid DNA transfection complexes for 48 h; or siRNA transfection complexes for 48 h (for viability assays or multi-cycle virus growth assays with influenza viruses in A459 cells), 60 h (for multi-cycle virus growth assays with WNV in A549 cells), or 72 h (for viability assays or multi-cycle growth assays with Ebola-ΔVP30 or MERS-CoV in Huh7 cells). For siRNA experiments, we verified high transfection efficiency by visually comparing cells treated with non-targeting and transfection efficiency controls prior to proceeding with further analyses.

### Viability assay

To determine how HB gene perturbation affects cellular viability, we assayed total intracellular ATP levels in A549, 293T, or Huh7 cells by using the CellTiter-Glo kit (Promega Corp.) and a Tecan plate reader.

### Quantitative real-time PCR

To confirm siRNA-mediated HB gene knockdown, we used quantitative real-time PCR (qRT-PCR) to measure target mRNA levels (in triplicate) in total RNA extracted from A549 cells by using the RNeasy Mini kit (Qiagen). We performed first-strand cDNA synthesis with the QuantiTect reverse transcription kit (Qiagen); carried out qRT-PCR reactions with gene-specific SYBR green-based primer assays (either QuantiTect assays from Qiagen or PrimeTime assays from IDT, Inc.) and PowerUp SYBR green master mix (Applied Biosystems, Thermo Fisher Scientific Corp); and quantified mRNA levels with the QuantStudio 6 Flex Real-Time PCR System (Applied Biosystems). For each high-ranked and low-ranked HB candidate gene, we determined relative mRNA quantities by using the comparative threshold cycle (ΔΔCt) method, with the GAPDH gene serving as the endogenous reference and RNA from non-targeting control siRNA-treated cells serving as the calibrators. A list of all qRT-PCR primer assays (including the primer assay types and product numbers) is provided in **Supplementary Table 5c**.

### Immunoblotting

To confirm cDNA overexpression, we used immunobloting to assess HB protein levels in total proteins extracted from 293T cells by using NP40 lysis buffer (10 mM Tris-HCl [pH 7.5], 150 mM NaCl, 1 mM EDTA, and 1% Igepal CA-630) containing a 1X protease inhibitor cocktail (cOmplete Mini EDTA-Free, Roche). We separated proteins by sodium dodecyl sulfate (SDS) polyacrylamide gel electrophoresis (PAGE), transferred them to polyvinylidene fluoride (PVDF) membranes (Invitrogen, Thermo Fisher Scientific Corp.), and performed blotting with a monoclonal anti-Flag M2 (cat. no. F1804, Sigma-Aldrich) or polyclonal anti-NFE2L3 (cat. no. ab11136, Abcam) primary antibodies, and horseradish peroxidase-conjugated secondary antibodies (Invitrogen). Immunoblots were developed with SuperSignal West Femto substrate (Thermo Fisher Scientific Corp.) and an AlphaImager (Alpha Innotech).

### Multi-cycle virus growth assays

After washing siRNA- or plasmid-transfected cells twice with phosphate-buffered saline (PBS), we inoculated cells with fresh medium containing a moi of 0.001 pfu/ffu per cell (pH1N1 and H3N2 influenza viruses, MERS-CoV, and WNV), 0.0001 pfu/ffu per cell (H5N1 influenza virus and Ebola-ΔVP30) and collected supernatants for virus quantification at 24 h (WNV and H5N1), 48 h (pH1N1, H3N2, and H5N1), or 72 h (Ebola-ΔVP30 and MERS-CoV) post-infection. Virus growth assays in A549 cells were performed in 1:1 DMEM/F12 medium supplemented with 0.6% bovine serum albumin fraction V (Sigma-Aldrich) and 0.2 µg/ml of N-tosyl-L-phenylalanine chloromethyl ketone (TPCK)-treated trypsin (Worthington Biochemical Corp.) (influenza viruses); or DMEM containing 2% FBS (WNV). Virus growth assays in Huh7 cells were carried out in the same media used for cell propagation (Ebola-ΔVP30 and MERS-CoV). To quantify the effects of HB gene perturbation on virus growth, we used standard pfu or ffu assays, as appropriate.

### Luciferase reporter assays

After washing siRNA- or plasmid-transfected cells once with PBS, we stimulated reporter gene expression as described below, incubated cells for 24 h, and then measured luciferase activity in cell lysates by using the Steady-Glo luciferase assay system (Promega Corp.) and a Tecan plate reader. Cells were stimulated as follows (in all cases, hygromycin was excluded from stimulation media): (*i*) For A549-IFNB1-Luc cells treated with siRNAs or 293T cells transfected with plasmids, we stimulated type I IFN expression by either infecting with Sendai virus (moi = 10 pfu per cell) or transfecting with 1 µg per well of polyinosine-polycytidylic acid (poly(I:C), Sigma-Aldrich) via the LT-1 transfection reagent (Mirus Bio LLC) in A549 or 293T cell propagation medium; (*ii*) For A549-NFκB-Luc cells treated with siRNAs or 293T cells transfected with plasmids, we stimulated NFκB-dependent gene expression by treating cells with purified human TNF (100 ng per ml; Cell Signaling Technology, Inc.) in 1X Opti-MEM reduced serum media (Gibco Life Sciences) containing 0.25% BSA fraction V; and (*iii*) For 293T cells transfected with plasmids, we stimulated ISG expression by treating cells with purified human IFNB1 (1,000 U per ml; PBL Assay Science) in 1X Opti-MEM containing 0.25% BSA fraction V. For all siRNA transfection experiments, we also carried out mock-stimulations. Mock stimulation protocols included treatment with fresh A549 growth medium lacking Sendai virus, transfection with LT-1 transfection reagent lacking poly(I:C), or treatment with Opti-MEM lacking purified human IFNB1 or TNF.

### *Nfe2l3* knockout mouse breeding

*Nfe2l3* knockout (KO) mice were generated previously^52^. Briefly, they were derived from 129S6 embryonic cells, backcrossed to C57BL/6J mice (strain #000664, The Jackson Laboratory) nine times, inbred for 8 generations, backcrossed to C57BL/6J three more times, inbred for 2 generations, and back-crossed to C57BL/6J one final time. We used heterozygous breeder pairs (*Nfe2l3*(+/–)) from the final backcross to establish a breeding colony for the current study. For five progeny mice generated from the breeder pairs, we carried out genome scanning analysis with two unique single nucleotide polymorphism (SNP) panels (The Jackson Laboratory) that detect either 150 SNPs polymorphic between C57BL/6 and 129S1/SvIm strains (panel 1) or 150 SNPs polymorphic between C57BL/6J and C57BL/6NJ strains (panel 2). Progeny mice were completely homogenous, possessing 100% C57BL/6 SNPs (panel 1) and 100% C57BL/6J SNPs (panel 2). Therefore, we inbred the mice over four additional generations to produce WT and KO littermates for influenza virus pathogenicity studies. Subsequently, we carried out one additional backcross to C57BL/6J mice, and heterozygous progeny were inbred for two more generations to produce WT and KO littermates for tissue collection experiments.

### *Nfe2l3* genotyping

Genomic DNA was released from mouse tail biopsies by incubation in DirectPCR Lysis Reagent (Tail) (Viagen Biotech, Inc.) at 85°C, and used directly for standard PCR reactions (50 µl total volume), including 2 µl of the lysed genomic DNA template, 5% DMSO, and three primers: M2 5’HOM-F (5’ CCAGACCAGGTTTGGCTTGGT), M2 3’HOM-R (5’ GGGTCACCACAGACTAGTACT), and M2 9512-R (5’ TGGGATGGGGTGTTAAGAGA). Cycling conditions were as follows: (***i***) 94°C for 2’; (***ii***) 30 cycles of 94°C for 30”, 62°C for 30”, and 72° for 1’; and (***iii***) 72°C for 10’. PCR amplicons (WT ∼460 bp and KO ∼320 bp) were evaluated by 0.7% agarose gel electrophoresis.

### Mouse infections and sample collection

We anesthetized groups of 8-10-week-old female WT and KO mice by intraperitoneal (i.p.) injection of ketamine and dexmedetomidine (45-75 mg/kg ketamine + 0.25-1 mg/kg dexmedetomidine), performed intranasal inoculation with 50 µl of PBS (‘mock’) or PBS containing serial dilutions of OK38L3, and then reversed dexmedetomidine by i.p. injection of atipamezole (0.1-1 mg/kg). We monitored individual mouse body weights and survival daily for up to 16 days, and humanely euthanized mice at the end of the observation period, at designated time points (for tissue collection) or when mice exhibited severe clinical symptoms. We obtained nasal turbinate and lung tissues after euthanizing mice and froze all tissue portions at -80°C in the absence of buffer until further analysis.

### Virus titration in mouse tissues

For virus titration, we thawed and weighed nasal turbinates or lung tissue portions (right superior lobes), and then homogenized tissues in 1 ml PBS containing a penicillin/streptomycin mixture by using a TissueLyser II (Qiagen) at 30-Hz oscillation frequency for 3 min. We centrifuged homogenates to remove debris (10,000 x *g* for 10 minutes), quantified virus titers (pfu) in clarified homogenate supernatants by using standard plaque assays in MDCK cells, and normalized the titers to pfu per gram of nasal turbinate or lung tissue.

### Cytokine quantification in lung tissues

For cytokine quantification, we prepared clarified homogenate supernatants of lung tissues (right inferior lobes) as for virus titration, except that the homogenization buffer included a 1X protease inhibitor cocktail (cOmplete Mini EDTA-free, Roche) and the volume was 500 µl. We quantified cytokine concentrations with the Bio-Plex Pro Mouse Cytokine 23-plex Assay and the Bio-Plex 200 system (Bio-Rad Laboratories, Inc.), or with VeriKine Mouse Interferon Alpha or Beta ELISA kits (PBL Assay Science) and a Tecan plate reader. Cytokine concentrations were normalized to picograms (pg) per gram of lung tissue.

### Statistical analyses

Enrichment *p*-values for biological pathway and process terms were determined by the Metascape web tool^27^ and transformed to (—) log *p*-value enrichment scores for graphical representation. All other statistical tests were performed by using GraphPad Prism version 8.0.0 for Windows (GraphPad Software). To compare the means of pH1N1 or H5N1 virus titers in negative control and NP siRNA treated cells, we log_10_-transformed virus titer values and used unpaired, two-tailed t-tests with a Welch’s correction. For siRNA experiments assessing the effect of HB candidate gene knockdown on multi-cycle virus growth or host pathway activation (via promoter-reporter assays), we compared the means of log_10_-transformed virus titers or luciferase values (relative light units, RLU) between replicates of individual test conditions (*i.e.*, cells perturbed by an siRNA targeting a HB gene mRNA) and batch-specific negative controls by using unpaired, two-tailed t-tests, followed by a Holm-Sidak post-test to calculate multiplicity-adjusted *p*-values. For graphical representation, we calculated log_2_–transformed virus titer or luciferase RLU ratios for individual test siRNAs (*i.e.*, mean test siRNA titer or luciferase values versus mean negative control siRNA titer or luciferase values). To determine whether the hit rate of high-ranked HB candidate genes differed from that of low-ranked HB candidates, we generated a contingency table and compared the distribution of hits and non-hits in pH1N1 infections by using a one-sided Fisher’s exact test.

For cDNA experiments assessing the effect of HB candidate gene overexpression on multi-cycle H5N1 virus growth, we compared the means of log_10_-transformed virus titers between replicates of individual test conditions (*i.e.*, cells transfected with a plasmid expressing an HB cDNA) and the time-matched negative control by using ordinary one-way analysis of variance (ANOVA). For cDNA experiments assessing the effect of HB candidate gene overexpression on host pathway activation, we combined data from two or three biological replicate experiments, each with three technical replicates. First, within each experiment, we normalized each replicate luciferase activity value by dividing by the mean luciferase activity value of the negative controls. Then, we log_10_-transformed and aggregated the normalized replicate values for each test condition across replicate experiments and compared the means of test and negative control conditions by using ordinary one-way ANOVA with a multiplicity correction.

To compare body weights of *Nfe2l3* WT and KO mice infected with pH1N1, we log_10_-transformed weight ratios for each mouse at each time point, and then compared group weight loss profiles by using a mixed effects model with a Geisser-Greenhouse correction. This analysis generated a *p*-value for the overall effects of *Nfe2l3* expression on mouse body weight loss, as well as *p*-values for effects observed at each time point. In WT and KO mice, we compared means of log_10_-transformed virus titers in nasal turbinate or lung tissues by using unpaired, two-tailed t-tests; and means of log_10_-transformed cytokine concentrations by using unpaired, two-tailed t-tests followed by a Holm-Sidak post-test.

### Data availability

Transcriptomics datasets were previously reported^21, 48, 74^ and deposited in the National Center for Biotechnology Information (NCBI) Gene Expression Omnibus (GEO; https://www.ncbi.nlm.nih.gov/geo/) (accession numbers GSE37571 and GSE28166). All processed datasets generated in this study are reported in **Supplementary Tables**, and associated raw data are available upon request.

### Biosafety

All work with influenza viruses, Sendai virus, and the biologically-contained Ebola-ΔVP30 virus was performed at the University of Wisconsin (UW)-Madison. *In vitro* experiments with pH1N1 or H3N2 influenza viruses or Sendai virus were performed in biosafety level 2+ (BSL-2+) containment; *in vitro* experiments with H5N1 influenza virus were performed in an animal-enhanced biosafety level 3+ (ABSL-3+) containment laboratory; and *in vivo* experiments with pH1N1 were performed in an animal-enhanced BSL-2 (ABSL-2) laboratory.

Experiments involving the biologically-contained Ebola-ΔVP30 virus were carried out in BSL-2+ containment, under approval by the UW-Madison Institutional Biosafety Committee, the US CDC, and the US National Institutes of Health. *In vitro* experiments with MERS-CoV or WNV were performed in BSL-3 containment laboratories at the University of North Carolina at Chapel Hill (UNC-Chapel Hill) or Washington University in St. Louis (WUSTL), respectively. The US CDC and/or the US Department of Agriculture approved the use of BSL-2+, ABSL-2, BSL-3, and ABSL-3+ containment facilities at the UW-Madison, the UNC-Chapel Hill, and WUSTL.

### Ethics statement

All animal experiments and procedures were approved by the UW-Madison School of Veterinary Medicine Animal Care and Use Committee (protocol # V006426-A04) under relevant institutional and American Veterinary Association guidelines.

## ACKNOWLEDGEMENTS

This study was funded by grant U19AI106772 from the National Institute of Allergy and Infectious Diseases, National Institutes of Health (USA); grants JP18am001007, JP18fm0108006, JP17fk0108202, JP18fk0108104, JP19fm0108006, JP22wm0125002, and JP22fk0108626 from the Japan Agency for Medical Research and Development; and grants JP16H06429, JP16K21723, and JP16H06434 from the Ministry of Education, Culture, Science, Sports, and Technology of Japan. Pacific Northwest National Laboratory is a multi-program laboratory operated by Battelle for the U.S. Department of Energy under Contract DE-AC05-76RL01830. The authors would like to thank Zachary Najacht and Daniel Beechler for technical assistance.

## SUPPLEMENTARY FIGURE LEGENDS

**Supplementary Figure 1.**
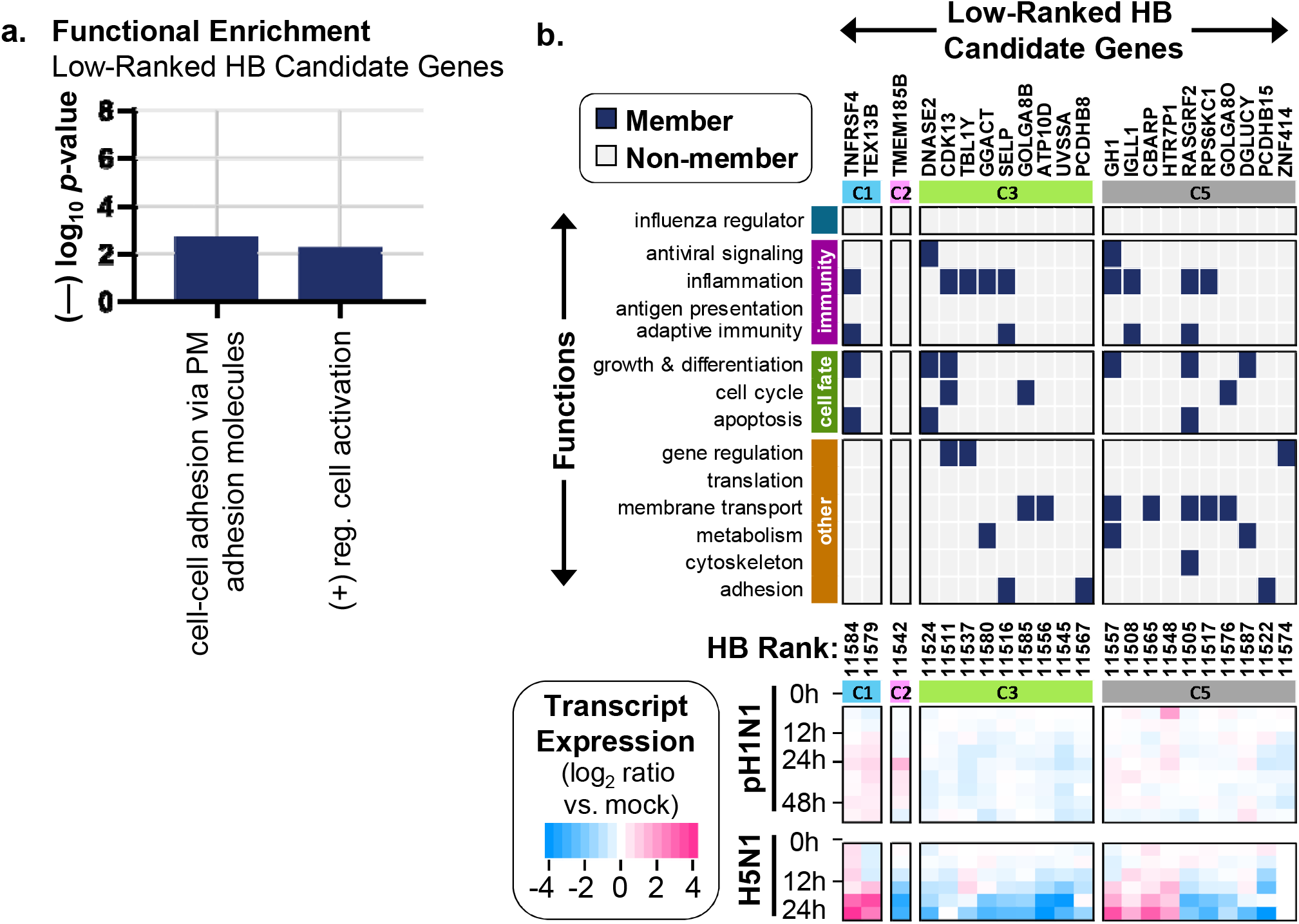
Attributes of low-ranked HB genes. **(a)** For 22 low-ranked HB genes (**Supplementary Table 1b**), the graph shows representative enriched functional terms and enrichment scores (—log_10_ *p*-values) determined by the web-based Metascape software (also see **Supplementary Table 1d**). **(b)** The panel shows individual functions of low-ranked HB genes (if known) (**top**) and their expression in Calu-3 cells infected with pH1N1 (A/California/04/2009) or H5N1 (A/Vietnam/1203/2004) (**bottom**) (also see **Supplementary Table 1f**). HB gene expression clusters (C1, C2, C3, and C5; the same as those described for high-ranked HB genes (Fig. 1c)) are indicated above both the function and expression sub-panels, HB gene ranks (see **Supplementary Table 1a**) are shown between the sub-panels, and a heat map key is given at the bottom left.

**Supplementary Figure 2.**
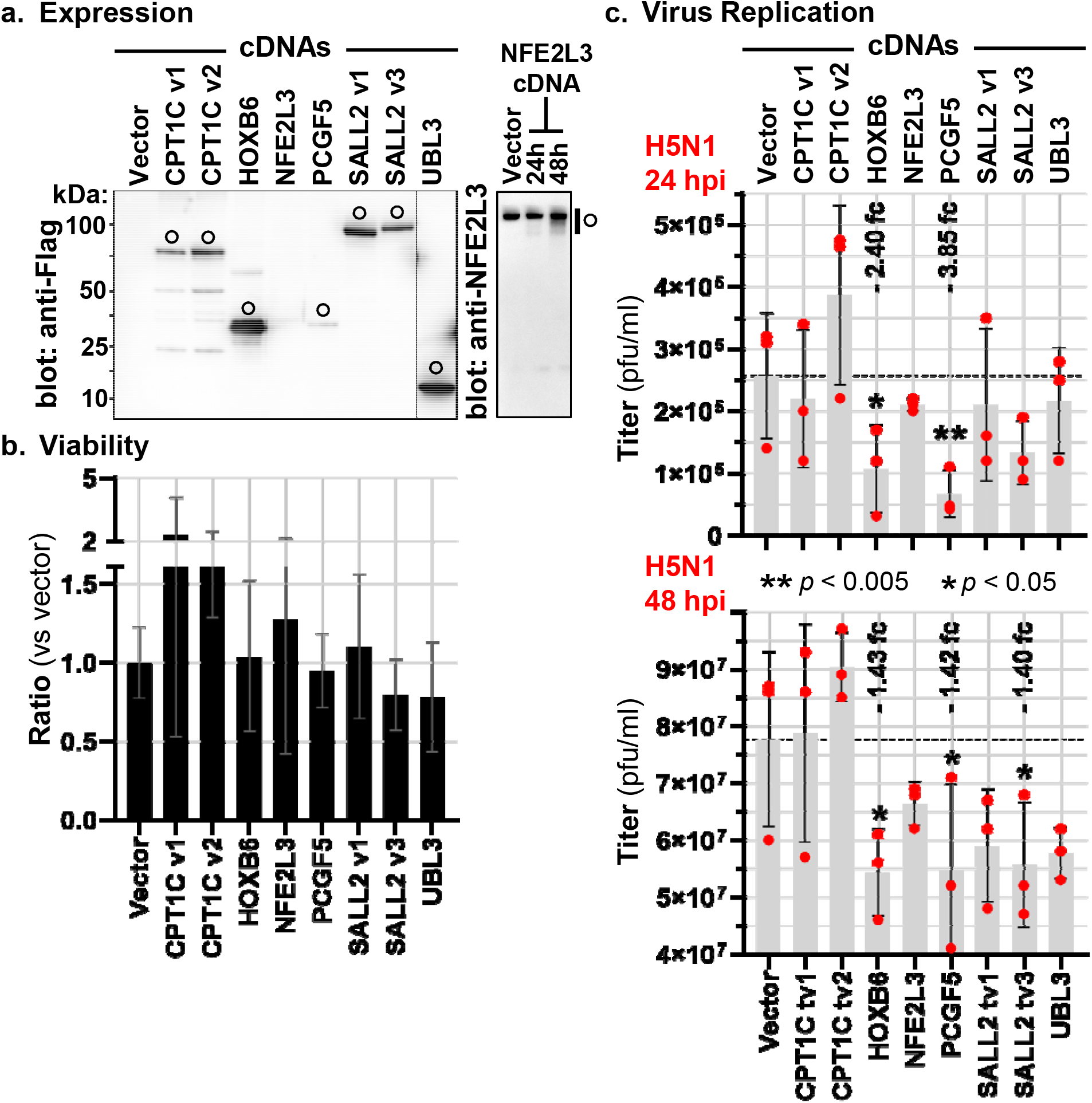
H5N1 virus growth in cells transiently overexpressing high-ranked HB genes with antiviral activity against pH1N1. (**a**) The panel depicts representative immunoblots of 293T cell lysates after transfection with an empty vector or plasmids expressing HB gene cDNAs carrying the Flag epitope tag (**left**) or lacking any epitope tag (**right**). Open circles indicate the full-length overexpressed protein for each HB gene (estimated molecular weights are provided in **Supplementary Table 5b**). Comparable results were observed in 4 other independent experiments. (**b**) The panel shows the mean viability ratios of 293T cells transfected with plasmids expressing HB gene cDNAs or an empty vector (negative control). We used the luciferase-based Cell Titer Glo assay (Promega) to measure intracellular ATP levels and calculated ratios for each HB cDNA versus the empty vector. The plotted data represent pooled data from 3 independent experiments, each with 3 replicates per cDNA. (**c**) 293T cells transfected with plasmids expressing HB genes were inoculated with influenza A/Vietnam/1203/2004 (H5N1) at a multiplicity of infection of 0.0001, and supernatants were collected for virus titration by plaque assay at 24 h (**top**) or 48 h (**bottom**) post-infection. In both plots, individual replicate titer values (3 per cDNA at each time point) are represented by the red dots, gray bars show the mean titer, and variation is represented by standard deviation. At each time point, mean titers of cells transfected with HB cDNAs were compared to that of empty vector transfections by using ordinary one-way ANOVA, and significant *p*-values are indicated on the figure panels. All data shown in panel (**c**) were collected in one experiment. fc, fold change.

**Supplementary Figure 3.**
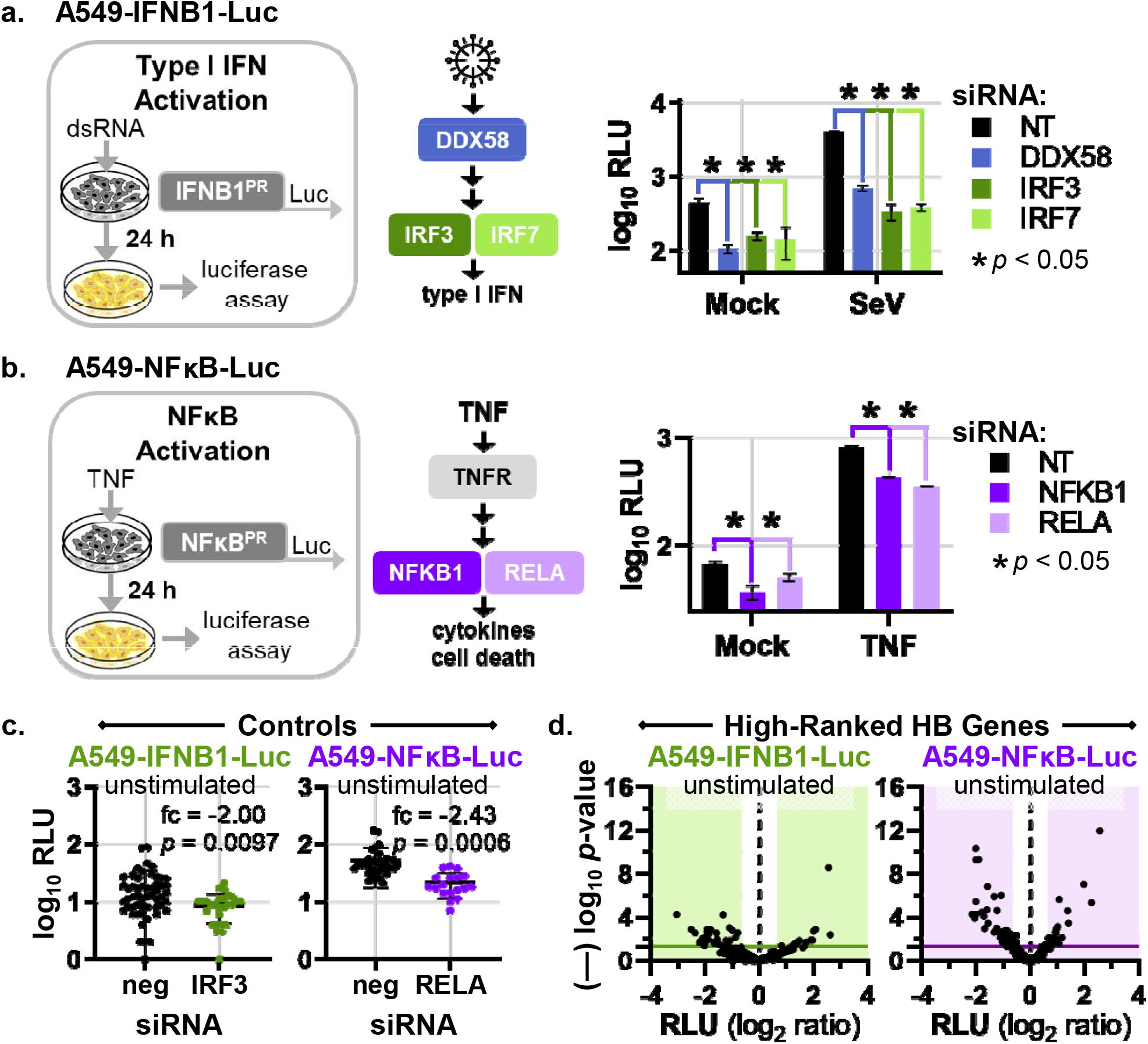
Generation and validation of type I IFN and NFκB promoter reporter cell lines. (**a**) and (**b**) depict generation and validation of A549-IFNB1-Luc or A549-NFκB-Luc cell lines, respectively. In each panel, the left-most segment shows a conceptual overview of the promoter (PR)-reporter cassette and a simple flow chart of promoter stimulation and activation measurement. The graphs at the right show promoter activation levels (relative light units, RLU) in mock-stimulated or stimulated cell clones treated with non-targeting control siRNAs (black bars) or siRNAs targeting specific components of each signaling pathway (colored bars) (three replicates per siRNA treatment and stimulation condition). The data were collected in two independent experiments (one experiment per cell line). Mean RLU were compared by using unpaired, two-tailed Student’s t-tests and *p*-values are indicated on the graphs. (**c**) Graphs show RLU values in unstimulated A549-IFNB1-Luc (**left**) or A549-NFκB-Luc (**right**) cells treated with control siRNAs. Mean RLU values of cells treated with negative control siRNA (neg) and siRNA targeting either IRF3 or the RELA component of the NFκB transcription factor were compared by using two-tailed, unpaired t-tests, and the resultant fold changes and *p*-values are indicated on the graphs. The data in the left and right panels each represent 4 independent experiments. (**d**) Volcano scatter plots depict log_2_-transformed mean RLU ratios (*x*-axis) versus (—) log_10_ *p*-values (*y*-axis) for unstimulated reporter cells treated with siRNAs targeting 50 high-ranked HB genes (2-4 siRNAs per gene) and assayed (in triplicate) for effects on IFNB1 (**left**) or NFκB (**right**) activation (**Supplementary Table 3a and 3b**). To determine ratios, we compared mean RLU values for HB gene siRNA- and negative control siRNA-treated cells, and *p*-values were calculated by using by using two-tailed, unpaired t-tests. In each plot, a data point represents the outcome for a single HB gene siRNA, the shaded areas demarcate +/-1.6-fold change in RLU, and a solid horizontal line indicates *p* = 0.05. Each siRNA was evaluated in one experiment in each reporter cell line, and complete datasets were collected in a series of 4 independent experiments for each cell line and pooled to generate the plots.

## LIST OF SUPPLEMENTARY TABLES

- **Supplementary Table 1. HB gene ranks, expression, and functional enrichment.**
- **Supplementary Table 2. pH1N1 and H5N1 siRNA screening data and hit identification for high- and low-ranked HB genes.**
- **Supplementary Table 3. A549-IFNB1-Luc or A549-NF**κ**B-Luc screening data and hit identification for high-ranked HB genes.**
- **Supplementary Table 4. Other human virus siRNA screening data and hit identification for selected high-ranked HB genes.**
- **Supplementary Table 5. Reagents (siRNAs, qRT-PCR primers, and cDNAs).**

